# Transfer Learning and Permutation-Invariance improving Predicting Genome-wide, Cell-Specific and Directional Interventions Effects of Complex Systems

**DOI:** 10.1101/2025.04.07.647536

**Authors:** Boyang Wang, Pan Boyu, Tingyu Zhang, Qingyuan Liu, Shao Li

## Abstract

With the advent of precision medicine, single-drug treatments may not fully satisfy the pursuit of precision medicine. However, single-drug treatments have accumulated a large amount of data and a considerable number of deep learning models. In this context, using transfer learning to effectively leverage the existing vast amount of single-compound intervention effect data to build models that can accurately predict the intervention effects of complex systems is highly worth investigating. In this study, we used a deep model based on permutation-invariance as the core module, pre-trained on a large amount of single-compound intervention data in cell lines, and fine-tuned on a small amount of complex system (like natural products) intervention data in cell lines, resulting in a predictive model named SETComp (the Concat version with ~200M parameters and the Add version with ~173M parameters). The two versions of SETComp achieved an accuracy of 93.86% and 92.70%, respectively, on the complex system-cell-gene association test set, improving by 5.82% to 27.59% compared to the baseline. When predicting the intervention effects of those complex systems the model had never encountered before, the accuracy increased by up to 24.83% compared to the baseline. In our in vitro validation, up to 88.65% of the predictions were confirmed to be correct, and the model’s output showed a significant positive correlation with the real-world foldchange. We further observed SETComp’s potential in various biomedical scenarios, achieving good performance in applications such as mechanism uncovering, repositioning, and compound synergy discovery.

## Introduction

With the advancement of artificial intelligence and high-throughput omics technologies, researchers have gained a deeper understanding of life science or biomedical issues such as disease mechanisms and drug development. It has gradually become evident that single drugs may not adequately address the complex mechanisms underlying complex diseases. In this context, combination therapies involving multiple drugs or treatments based on complex systems like natural products (NP) have gradually become more popular, with predicting the targets of multiple drugs or complex systems with high throughput and high accuracy emerging as a hot topic in biomedicine. However, compared to target prediction for single compounds^1–5^, target prediction for complex systems requires more consideration on how to integrate compound–compound interactions, compound side effects and other information. Additionally, compared to target prediction for combinations of two or three compounds^6–8^, predicting targets for complex systems like NP, which involve an indeterminate number of compounds, appears more challenging. Currently, there are few existing algorithms for target prediction of complex systems, particularly network-based approaches. Li et al.^9^ proposed statistical modeling based on the targets of all compounds to predict the holistic targets of complex systems. Zhou et al.^10–12^ treated complex systems as whole entities and utilized heterogeneous graph neural networks for prediction. However, these prediction methods have certain limitations, including the inability to make predictions based on specific entities such as target cells, low prediction accuracy, no ability for directional prediction and a reliance on extensive prior knowledge, such as the need for the targets of compounds contained within NP. Therefore, developing a cell-specific target prediction model and a target promotion/inhibition direction prediction model suitable for complex systems with a large number of compounds is highly valuable and holds great potential for biomedical application.

Transfer learning, as a deep learning technique^13,14^, has excellent applicability in scenarios with limited sample sizes. Till now, there is a large amount of data accumulated on compound-intervened transcriptomics like Connectivity Map (CMap) for The Library of Integrated Network-Based Cellular Signatures (LINCS) project^15^. In comparison, transcriptomic data related to complex systems is relatively scarce in comparison. Therefore, applying transfer learning to complex systems-related applications is highly meaningful. Similarly, set-based deep learning possesses the property of permutation invariance, with Deep Sets^16^ and Set Transformer^17^ being classic models in this domain. Set-based deep learning methods are designed to handle unordered data, such as sets, by ensuring that the model’s output remains invariant regardless of the input data’s order. This property makes them particularly effective for tasks where the order of elements in the set is irrelevant, such as in complex systems with multiple compounds. These models provide a powerful approach to understanding and predicting the interactions and effects of various components within these systems.

In this study, we employed transfer learning techniques by treating complex systems as sets of compound combinations and utilizing permutation-invariant (set-based) deep learning as the core model framework, named Set Embedding and Transfer learning model for Complex systems (SETComp), to predict genome-wide, cell-specific and directional targets for both compounds and complex systems. We pre-trained the model on 970,481,750 compound–cell–gene association data obtained from transcriptomics data processed from LINCS and further fine-tuned it on 2,579,488 natural product–cell–gene association data collected and processed from GEO datasets and literature. Two versions of SETComp (Concat version with ~200M parameters and Add version with ~173M parameters) achieved 93.86% and 92.70% accuracy in predicting NP-cell-gene associations, respectively, of which the performance was also tested in real-world in vitro assays, achieving high accuracy. Besides, in multiple downstream application scenarios, SETComp has demonstrated good performance, including revealing the potential mechanisms of action of complex systems on different cell lines corresponding to tumors, repositioning complex systems to discover new potential target diseases, and predicting the gene-level synergistic interactions of complex systems and their constituent compounds. In summary, we have trained a model based on set-based deep learning using vast amounts of compound-cell-gene data and applied transfer learning to achieve high-precision predictions of complex system-cell-gene associations, which demonstrates strong application potential in various downstream scenarios related to biomedical issues.

## Result

### Overview of the study

In this study, we proposed a model named Set Embedding and Transfer learning model for Complex systems (SETComp), cored by the combination of transfer learning and set-based deep learning. Aiming at predicting transcriptome-level targets in different types of cells, SETComp was designed for stimulating the genome-wide changes in the cellular level after the intervening of complex compound systems like drug combinations or NP (Figure 1a).

**Figure 1.**
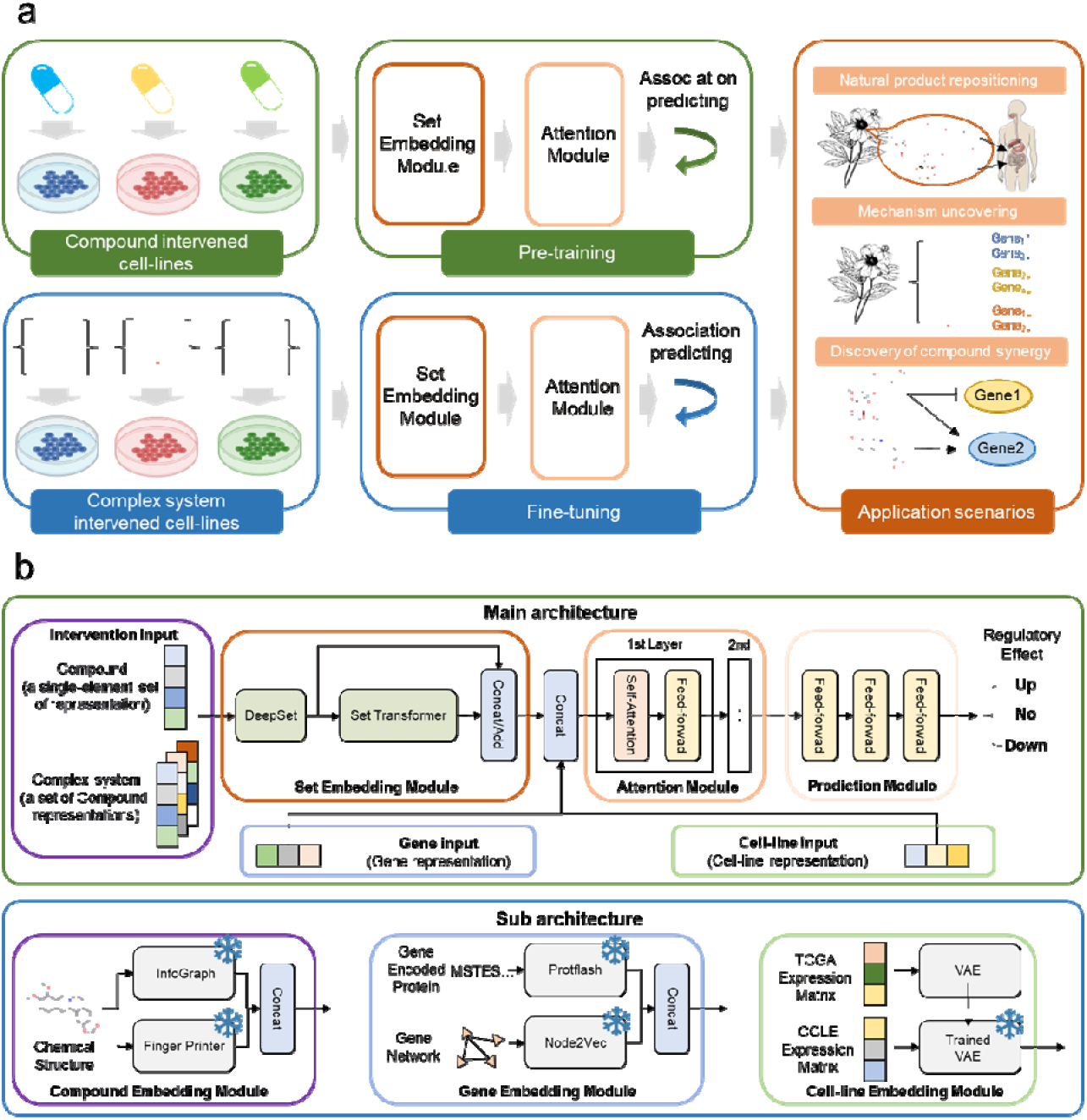
Model schematic of the SETComp model. (a) The workflow of SETComp. The model is pre-trained on compound-cell-gene association extracted from CMap in LINCS project, and fine-tuned on complex system (like natural products)-cell-gene association processed from collected GEO datasets and literature. The fine-tuned SETComp model demonstrates scalability across multiple biomedical-related application scenarios, including repositioning, mechanism uncovering, and discovery of compound synergy. (b) The architecture of SETComp. The core module of SETComp lied in the permutation-invariance Set Embedding Module, which permutes-invariantly characterizes the complex system packaged as a set (with initial characterization through Deep Sets and further deep characterization through both Deep Sets and Set Transformer, combined at the end of the module in a concatenation/addition form). The set-based embedding is then combined with gene representations and cell line representations, followed by a self-attention module to learn the relationships between features. Finally, the prediction module, consisting of an MLP, performs association classification, including upregulation, downregulation, and no significant association. The compounds are jointly represented by the self-supervised pre-trained InfoGraph model and the Finger printer model, while genes are represented by a combination of PPI network-based node2vec and the pre-trained protein language model Protflash. Cell lines are represented based on the latent layer of the trained VAE. During feature extraction, all models in these sub modules are frozen.

To achieve modeling the effects at the transcriptomic level after the intervention of a single compound or complex systems like NP in different cell conditions (Fig S1), SETComp consists of three frozen encoders and three internal modules for feature exacting, feature integration and performing prediction (Figure 1b). The three frozen encoders are: a compound encoder, which combines the trained Infograph graph neural network and a Fingerprinter encoder; a gene encoder, integrating a lightweight pre-trained large language model (LLM) with a node2vec model derived from protein-protein interaction (PPI) networks; and a cell state encoder, which is a Variational Auto-Encoders (VAE) model trained on expression profiles from The Cancer Genome Atlas (TCGA). As for the internal modules, centered around the set embedding module, which consists of an initial embedding based on Deep Sets^16^ and a further embedding based on Set Transformer^17^, the three internal modules also include a self-supervised module and a prediction module composed of multiple progressively dimension-reduced Multilayer Perceptron (MLP).

Compounds with SMILES description, genes, and cell lines were embedded into feature space with the three frozen encoders, respectively. SETComp was firstly pre-trained on 970,481,750 compound–cell–gene association data obtained from 1,805,898 transcriptomics samples with a single intervention of 39,321 compounds, to achieve accurate prediction of transcriptomics-level changes. With the collected relationships between compounds and NP, every natural product was considered as a set of compounds, and the model was then fine-tuned on 2,579,488 natural product-cell-gene pairs extracted from collected data obtained from both the GEO database (update to 2024 May) and literature for predicting transcriptomics-level targets for complex compound systems like NP. In addition, in vitro experimental validations were conducted on different cell lines and NPs to estimate the performance of SETComp in real-world assays. In the downstream, SETComp has been tested for applications in multiple biomedical-related scenarios, including the analysis of intervention mechanisms at the molecular and pathway levels, drug repositioning, and the analysis of synergistic effects between multiple compound components in complex systems.

### Pre-training of the SETComp model

Although RNA-seq technology has been around for over a decade and has accumulated a large amount of data in the context of single compound interventions, such as the CMap from LINCS project, data in the case of complex system interventions, such as NP interventions, is relatively lacking compared to single compounds. We have collected transcriptomic data from compound interventions in CMap which then enhanced by a modified CycleGAN^18^, transcriptomic data from NP interventions from GEO and literature (Fig 2a), original transcriptomics data from TCGA and CCLE, gene association networks and protein coding information from STRING^19^, compound structure information from PubChem^20^, and compound composition information for NP from HERB (v2.0)^21^ (Fig S2a). After cleaning, screening, enhancement, and preprocessing the transcriptomic data of single compound interventions, we obtained gene differential expression data for over 20,000 compounds (Fig S2b) across more than 90 cell lines (Fig S2c) and constructed 970,481,750 compound-cell line-gene association pairs.

**Figure 2.**
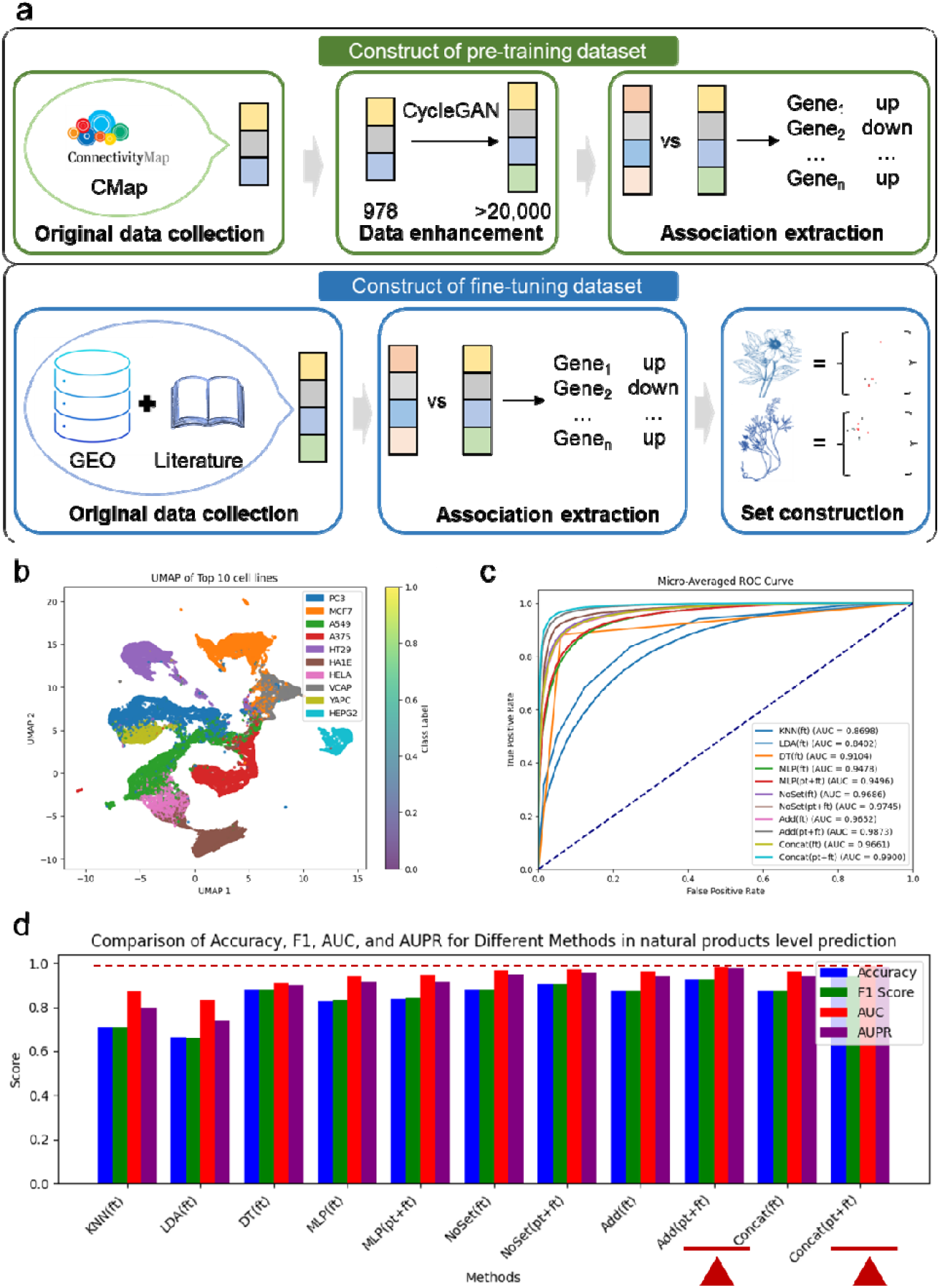
Preprocessing and performance of the SETComp model. (a) Illustration of the dataset construction of SETComp. The compound intervened expression profiles on multiple cell lines for pre-training were obtained from CMap in LINCS project, which were further cleaned and enhanced by a modified CycleGAN model. The compound-cell-gene associations were extract based on the differential expression analysis of the enhanced expression profile. The complex system (like natural products) intervened expression profiles on multiple cell lines for fine-tuning were obtained from GEO datasets and literature. And the complex system-cell-gene associations were also processed from the differential expression analysis. The sets of every complex system were constructed according to the recorded composition information from HERB (v2.0). (b) Detailed view of the embeddings of cell lines with largest number of intervened profiles involved in the pre-training data. After PCA dimensionality reduction of the original expression profiles of the top 10 cell lines with the most interventions, the data were further reduced using UMAP and visualized in a 2-dimensional space. (c) ROC curves of fine-tuned SETComp and baseline models on the test set. The baseline models include traditional machine learning models such as K-Nearest Neighbors (KNN), Linear Discriminant Analysis (LDA), and Decision Trees (DT), as well as a vanilla neural network (MLP) and the NoSet version consisting solely of the attention module and prediction module. (d) The performance of fine-tuned SETComp and baseline models on the test set. In the comparison of accuracy, F1 score, AUC, and AUPR, both the Concat version and Add version of the SETComp model achieved the highest levels, surpassing traditional machine learning models, vanilla neural network, and the NoSet version.

The enhanced transcriptomic expression data of untreated cell lines showed a discernible data distribution in UMAP space, indicating that the data-enhanced cell lines possess improved representational characteristics, better clustering, and more robust differentiation in gene expression patterns (Fig 2b, S2d). The core module of the SETComp model, the Set embedding module (Fig S3), consisted of two set-based deep learning model, Deep Sets^16^ and Set Transformer^17^ (see supplementary materials). Deep sets initially characterize the input sets (including single-element sets composed of individual compounds and multi-element sets composed of complex systems) as fixed-length sets, providing a preliminary representation of the input. The output after Deep sets representation is then used as a new set input, which is fed into the set Transformer. Through its unique Multihead Attention Block (MAB), Set Attention Block (SAB), Induced Set Attention Block (ISAB), and Pooling by Multihead Attention (PMA) blocks, further deep representation is obtained. Meanwhile, we employ two strategies, concatenation and addition, to combine the shallow set-based representation from Deep sets and the deep set-based representation obtained from the collaborative characterization of Deep sets and the set Transformer by either concatenating or adding them, to construct the Concat version (~200M parameters) and the Add version (~173M parameters) of the SETComp model. After concatenating the output of the set embedding module with the gene features and the cell line features obtained from the VAE, self-attention is applied to further learn the relationships between the features. Finally, the set-cell-gene prediction is performed through a prediction module composed of three layers of MLP with progressively reduced dimensions. After performing a grid search on the training parameters, including model parameters such as the size of the set Transformer, the size of MLP in the prediction module, as well as training parameters like batch size, learning rate, and regularization parameters such as L2 regularization and dropout, the optimal model parameters for the Concat version were used to determine the model structure (Table S1). Subsequently, the Add version also underwent a grid search on the training and regularization parameters (Table S2). Both versions of the SETComp model approach convergence after 5 epochs (Table S3, 4), and without overfitting, we trained both versions of the model to 10 epochs with the optimal parameters.

In our training set division for pre-training, all compounds belonging to NP components were extracted separately as the test set. By training on the entire training set, we found that the vanilla neural network composed only of the prediction module outperforms the two versions of the SETComp model (with AUCs of 0.9480 and 0.9417, respectively) in predicting compound-cell-gene associations, and further outperforms the model (the NoSet version), which consists of the self-attention module and the prediction module (Fig S4 a-d, Table S5). This may be because when predicting a single compound, treating it as an individual rather than as a set is more suitable and sufficiently saturated for the model. The two versions of the SETComp model perform better in predicting up-regulation and down-regulation, with an AUC of 0.96 for both (Fig S5 a-d). Further, on a smaller-scale training set (1/100 size of the full training set), we found that deep learning-based models performed better on the test set compared to traditional machine learning models (Fig S6 a-d, Table S6), including K-Nearest Neighbors (KNN), Linear Discriminant Analysis (LDA), and Decision Trees (DT).

### Fine-tuning improving the performance of prediction on complex systems

After completing the model pre-training, we focused on our research goal—achieving high-precision complex system-cell-gene association prediction. We pre-processed various complex system sets with compound features as elements, with the largest set containing up to 406 elements (Fig S7a). At the same time, we cleaned, pre-processed, and analyzed expression data of complex systems from the GEO database and literature across different cell lines (Fig S7b, c). We found that there was no significant correlation between the total expression levels of cell lines after complex system interventions, such as NP, and the number of compounds (Fig S7d) contained in NP (P=0.4331).

During the fine-tuning procedure, we also performed a grid search on the training parameters, including batch size and learning rate, as well as regularization parameters such as L2 regularization and dropout for the two versions of the SETComp model, to determine the optimal parameter combination (Table S7). We found that the performance of the two versions of the SETComp model, after pre-training and fine-tuning, surpassed traditional deep learning models such as KNN, LDA, and DT, as well as the vanilla neural network and the NoSet version in deep learning, and their performance when trained only on fine-tuning (Fig 2c). The two versions of the SETComp model achieved 93.86% and 92.70% in accuracy (Table S8), as well as 0.9888 and 0.9856 in AUC, respectively, improving the accuracy compared to the baseline machine learning model, increasing from 5.82% to 27.59%, while the AUC increases from 7.83% to 15.63% (Fig S8a, c) and the AUPR increases from 8.20% to 24.4% (Fig S8b). Compared to the vanilla neural network, the accuracy can be improved by 10.10%, and compared to the NoSet version, it shows an improvement of 5.74%.

Compared to the model that only underwent pre-training, we found that on the test set, the model that underwent both pre-training and fine-tuning performed better than the model that only underwent fine-tuning, and both outperformed the model that only underwent pre-training. This finding also demonstrates the benefit of fine-tuning on the prediction model, as well as the effectiveness of pre-training on compound-cell-gene in enhancing the model’s performance (Fig 3a, S9a-c, Table S9). At the same time, this finding also indicates that, beyond the superiority of deep learning itself in this task, the attention module and set embedding module progressively contribute to the improvement of the model’s performance. Then, based on the pre-trained model, we gradually added the size of the fine-tuning training set from 0% to 100%, and observed that as the size of the fine-tuning training set increased, the model’s performance on the fine-tuning test set gradually improved for both the Concat version (Fig 3b, S10a, b) and the Add version (Fig S10c) of the SETComp model. Furthermore, by adding models pre-trained on a small-scale pre-training dataset, we also performed fine-tuning and found that a larger pre-training dataset led to better performance on the fine-tuning test set, while accelerating the improvement of Accuracy and AUC (Fig 3c, S11c, d) as well as the reduction of loss (Fig S11a, b), and improving Accuracy and AUC of the converged models (Fig 3e, Fig S12a-d, Table S10).

**Figure 3.**
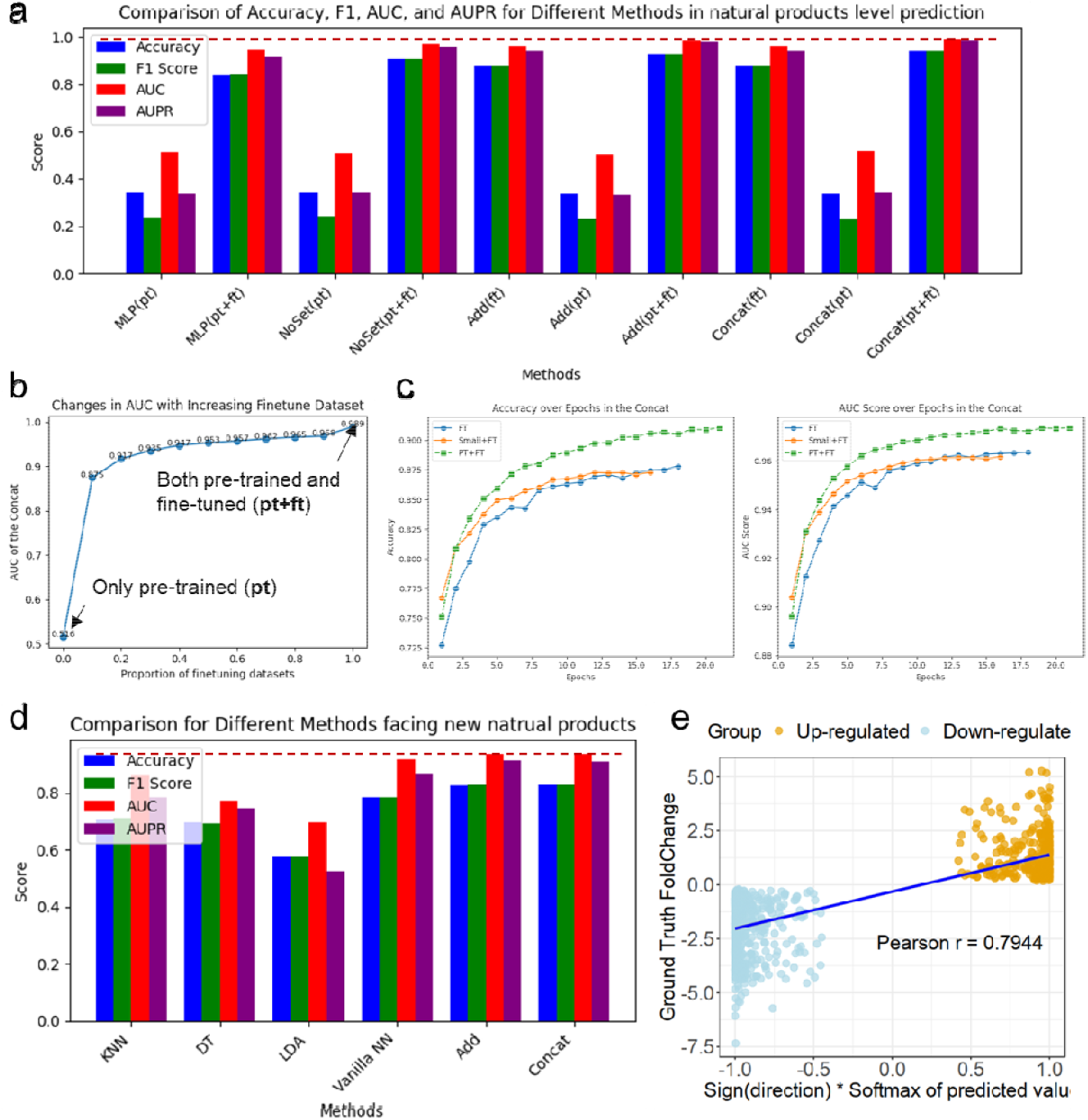
Ablation experiments and generalization study of the SETComp model. (a) The performance of fine-tuned SETComp and other deep learning models on the test set trained with different training datasets. In the comparison of accuracy, F1 score, AUC, and AUPR, both the Concat version and Add version of the SETComp model, after pre-training and fine-tuning, achieved the highest levels, surpassing the vanilla neural network and the NoSet version. They also outperformed their own performance when trained only on pre-training data or only on fine-tuning data. (b) A line chart showing how the AUC score of the Concat version of SETComp on the test set changes with the increasing percentage of the fine-tuning dataset in the total fine-tuning dataset. As the percentage of the fine-tuning dataset in the total fine-tuning dataset increases, i.e., with the increase in the size of the fine-tuning dataset, the model’s AUC on the test set also increases. (c) A line chart displaying how the Accuracy and AUC scores of the Concat version of SETComp on the test set change with variations in pre-training conditions. Compared to models pre-trained on a small-scale pre-training dataset (1/100) and models with no pre-training, the model fine-tuned and trained on the full pre-training dataset achieved higher Accuracy and AUC scores. (d) The performance of the model and baseline models on the test set extracted from intervention data from complex systems that the model had not encountered. In the comparison of accuracy, F1 score, AUC, and AUPR, both the Concat version and Add version of the SETComp model achieved the highest levels, surpassing traditional machine learning models and vanilla neural network. (e) The relationship between the model’s predicted output and the Foldchange in real-world data. After the softmax transformation, the model’s predicted outputs for class 0 (up-regulated) and class 1 (down-regulated) associations show a significant positive correlation with the corresponding real-world data’s Foldchange.

Furthermore, to assess the model’s generalization ability, we re-divided the fine-tuning dataset based on NP, using 10% of the NP-related transcriptomic data as the test set, and the remaining data as the training and test sets for fine-tuning the model, and re-trained the pre-trained model. When predicting NP that the model has not seen by the model before, it can achieve accuracies of 82.75% and 82.66% for the two versions of the model (Fig 3d, Table S11), respectively, which is an improvement of up to 24.83% in accuracy compared to the baseline machine learning models and 5.59% to the vanilla neural network (Fig S13a-c). Finally, to explore the potential for quantitative prediction in the future, we analyzed the model output values (predicted as down-regulated if negative) after applying softmax on all NP-cell-gene associations in the fine-tuning test set, and their actual statistical test logFoldChange (Fig 3e). We found a significant correlation, and this correlation varied across different NPs, ranging from 0.6734 to 0.9226 (Fig S14). This finding suggests that the model’s predicted results have a significant positive correlation with the actual transcriptomic logFoldChange, and it somewhat indicates the feasibility of extending the model to quantitative prediction.

### In vitro experiments validating the performance of the SETComp model

To further validate the predictive performance of the model, we conducted real-world transcriptomics assays to verify the model’s predictions. In multiple cell lines, we tested the effects of several NPs, including cinnamon, codonopsis, and astragalus (Fig 4a), and obtained their RNA-seq counts through transcriptome sequencing for differential expression analysis.

**Figure 4.**
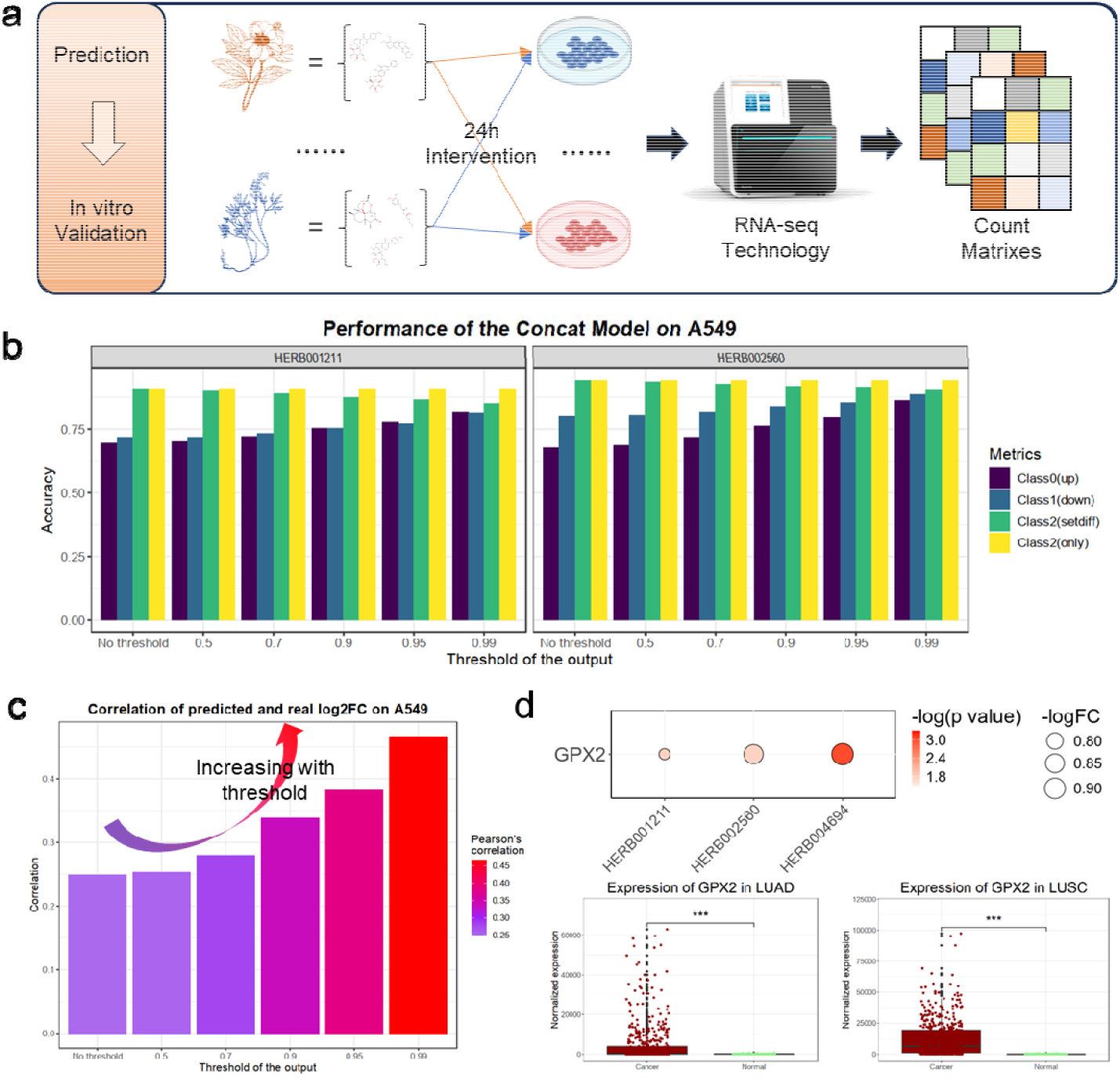
Case study of the SETComp model. (a) Case study illustration. We conducted 24-hour interventions of multiple complex systems (natural products) on cell lines and extracted RNA for RNA sequencing. (b) Accuracy of SETComp’s prediction of the intervention effects of Astragalus and Codonopsis in the A549 cell line within the differential expressed genes. In the A549 cell line, the accuracy of SETComp’s prediction of the intervention effects of Astragalus and Codonopsis for class 0 and 1, increases as the threshold increases, reaching a maximum of 88.65%. For the prediction of class 2, the accuracy can reach up to 94.01%. (c) The correlation coefficient between the SETComp predicted scores and the logFC obtained from real-world experiments within the differential expressed genes. As the threshold increases, the correlation coefficient between the SETComp predicted scores and the logFC from real-world experiments gradually increases, reaching a maximum of 0.4665. (d) Potential target GPX2 identified by combining model predictions, real transcriptomic data, and TCGA tumor data. After integrating the model’s predicted results (threshold 0.7) with transcriptomic intervention results, it was found that the three NPs collectively downregulated GPX2 and many other genes. Among them, six genes were found to be significantly overexpressed in tumor tissues of LUAD and LUSC. The upper bubble plot shows the downregulation of GPX2 in A549 cells after treatment with three NP in real-world transcriptomics assay.

After conventional bioinformatics processing (Fig S15a), we obtained the differential expression of each gene after cell line intervention. Among the differential genes (P value < 0.05), for the model-predicted upregulated targets of codonopsis in A549, 75.44% were found to be truly upregulated (with a threshold of 0.9), while 75.28% of the downregulated targets were confirmed to be downregulated; for the model-predicted upregulated targets of astragalus in A549, 76.25% were found to be truly upregulated (with the threshold of 0.9), and 83.95% of the downregulated targets were confirmed to be downregulated (Fig 4b). These accuracies improve as the threshold increases, reaching up to 81.72% for the upregulated prediction and 81.51% for the downregulated prediction for codonopsis at the threshold of 0.99; for astragalus, the upregulated prediction reaches 86.25% and the downregulated prediction reaches 88.65%. For cinnamon, the maximum accuracy in predicting downregulation can reach 83.12% (Fig S15b). Among all genes, in the prediction of upregulated genes, the accuracy under the intervention of codonopsis or astragalus can reach a maximum of 76.84% (Fig S15c), with accuracy showing an increasing trend as the threshold increases. In the prediction of negative samples (class 2), the model’s predictions remain relatively stable, with a maximum of 94.01% of predictions for class samples being confirmed as having no significant differences for astragalus, and 90.85% for codonopsis.

Similarly, we also observed the relationship between the scores predicted by the model for each gene under different NP interventions after softmax and the actual fold change of gene expression under each NP intervention. We found that the expression of intervention genes predicted by the model (classified as 0 or 1) is consistently significantly positively correlated with their true fold change, and as the threshold increases, the correlation score between them gradually increases (Fig 4c). In the differential genes of the A549 cell line, this correlation can reach a maximum of 0.4665, and in all genes, it can reach 0.2409 (Fig S16a). Focusing on the results of individual NP interventions, the positive correlation coefficient remains significant, and this phenomenon of increasing positive correlation with increasing threshold is still maintained. In the differential genes of the A549 cell line, the regularization coefficients under the interventions of astragalus, codonopsis, and cinnamon can reach 0.5788, 0.5486, and 0.4425, respectively (Fig S16b); in the full gene range of the A549 cell line, the regularization coefficients under the interventions of astragalus, codonopsis, and cinnamon can reach 0.2306, 0.2146, and 0.2778, respectively (Fig S16c). This finding further demonstrates that the model’s predictions not only enable qualitative directional predictions and predictions of intervention, but also possess the potential for quantitative prediction of relative expression levels, providing evidence support for subsequent quantitative predictive analyses in application scenarios.

Combining the prediction results of the SETComp model of multiple complex systems on A549, real-world transcriptomics assays and transcriptomics data from TCGA, we attempted to identify potential targets of some complex systems in the treatment of the A549-related clinical cancers, specifically non-small cell lung cancer LUAD and LUSC. Among the genes that were predicted by the model with high scores (threshold of 0.7), that genes showed significant differential expression in the transcriptome, and those were significantly different in both LUAD and LUSC, we found that some potential targets such as GPX2 (Fig 4d), PRR13, and APOC1, which are highly expressed in cancer samples of LUAD and LUSC (Fig S17b, c), were significantly downregulated under the intervention of all these three NP (Fig S17a). GPX2 has been studied in lung cancer, with reports indicating its involvement in apoptosis^22,23^, immune regulation^24^ and oxidative stress^25–27^, with clinical significance^28^ in lung cancer. The above findings, along with relevant literature, provide some evidence that SETComp’s predictive capability demonstrates strong generalization, robustness, and potential for application expansion.

### The extensive applications of the SETComp model in various biomedical issues

Last but not least, based on the model validated in both the test set and real-world assays, we use multiple biomedical application scenarios as downstream tasks to explore the practical application value of the model. First, we attempted to apply SETComp to explore the molecular and pathway mechanisms of complex system interventions in cell lines.

### Mechanism uncovering

Based on the aforementioned finding, that the model’s predicted values quantitatively reflect the activation/inhibition intensity of genes after intervention to some extent, we can apply Gene Set Enrichment Analysis (GSEA) to perform enrichment analysis on the prediction results and observe the potential pathway-level changes they may induce. We predicted the intervention effects of three NPs, including cinnamon, codonopsis, and astragalus (Fig S18a) on two cell lines, MCF-7 and A549, and constructed their potential regulatory molecular networks (Fig S18b-d). After GSEA based on the predicted up-regulated and down-regulated genes, astragalus was found to affect certain pathways in multiple modules in both MCF-7 and A549 cell lines (Fig 5a, b). Although astragalus potentially intervenes in pathways belonging to modules such as signal transduction, cell growth and death, and the immune system in both MCF-7 and A549 cell lines, the specific pathways within each module vary. For example, in cell growth and death, astragalus potentially regulates tumor cell growth and proliferation in MCF-7 by promoting the p53 signaling pathway^29–31^, while in the A549 cell line, it potentially promotes tumor cell apoptosis by enhancing the Apoptosis signaling pathway^32–34^. Additionally, in some classic signaling pathways, astragalus potentially inhibits tumor cell growth and proliferation in MCF-7 through the FOXO signaling pathway^35–37^, while in A549-related non-small cell lung cancer, it potentially alters the tumor immune microenvironment by promoting the Toll-like receptor signaling pathway^38,39^. Similar analysis also revealed that in the potential pathways of cinnamon and codonopsis interventions in MCF-7 (Fig S19a, b) and A549 (Fig S19a, b), the model is able to identify different potential molecules, pathways, and modular mechanisms for different complex systems intervening in different cell lines. Additionally, we observed the impact of different thresholds on the robustness of the analysis results. Taking the intervention of astragalus in MCF-7 and A549 as an example, we found that when the threshold was set to 0.7 (Fig S21a, b) or 0.9 (Fig S21c, d), the analysis results did not change significantly. The variations were likely observed in the same pathways but with different enrichment levels.

**Figure 5.**
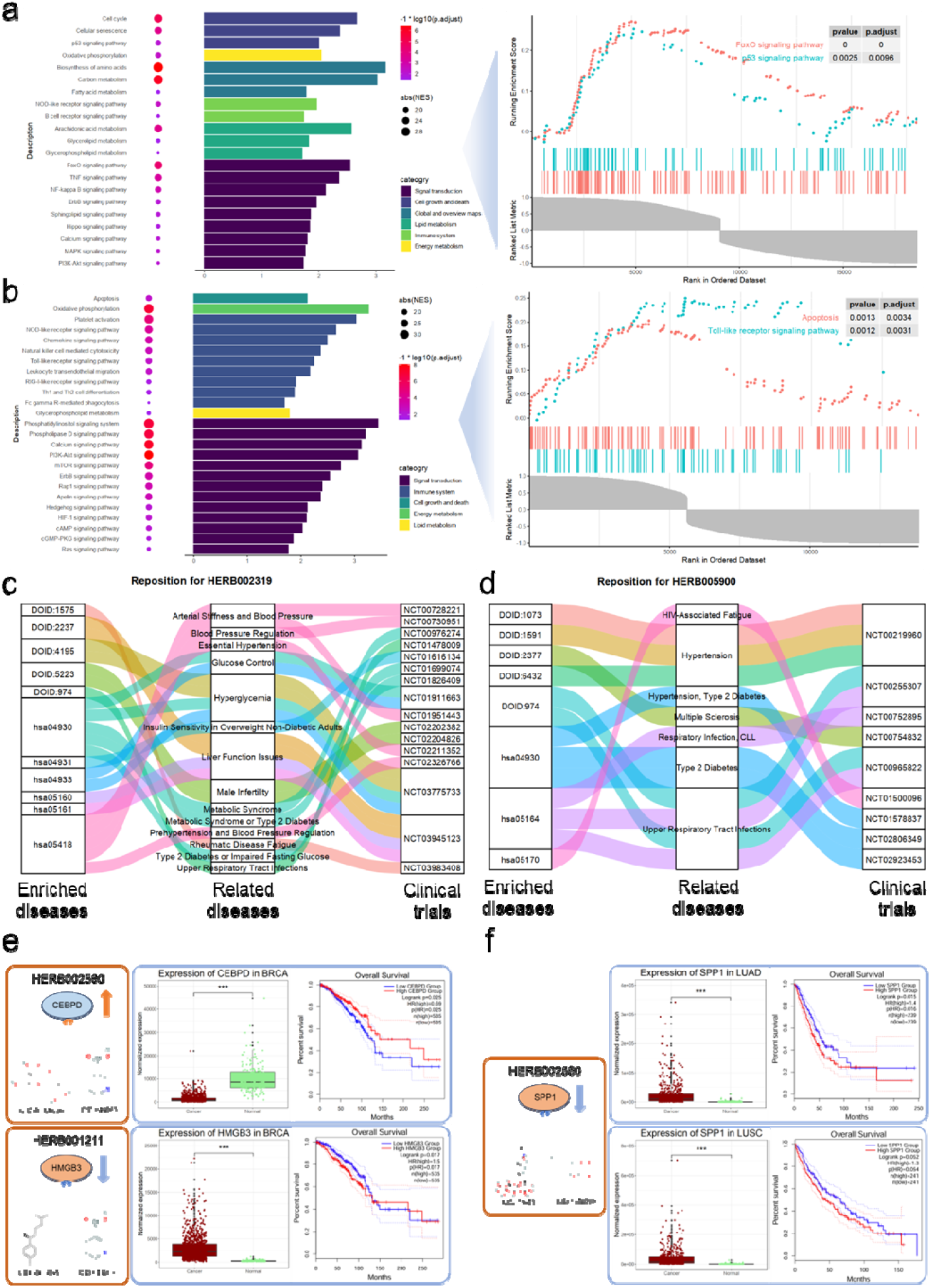
Application of SETComp in various biomedical scenarios. (a)-(b) Mechanism uncovering of potential intervention pathways of Astragalus in MCF-7 (a) and A549 cell lines (b) based on the prediction of SETComp. The GSEA enrichment analysis revealed that SETComp predicted Astragalus to be involved in multiple modules in both MCF-7 and A549 cell lines, like signal transduction, cell growth and death. However, the specific effects on MCF-7 and A549 cells differ. For example, Astragalus primarily affects the p53 signaling pathway in MCF-7, whereas in A549, it mainly potentially upregulates the apoptosis signaling pathway. (c)-(d) Reposition of certain complex systems like Red Ginseng (c) and American Ginseng (d) based on the prediction of SETComp. KEGG and DO enrichment analysis based on SETComp prediction results revealed that Red Ginseng and American Ginseng have potential interventions for multiple diseases, and this was supported by evidence from NCT clinical trials. (e) Discovery of compound synergy in complex systems intervening in BRCA based on SETComp. Based on SETComp’s predictions, multiple compounds in Astragalus (up) and Codonopsis (down) synergistically promote the expression of CEBPD and inhibit the expression of HMGB3 in MCF-7, respectively. It was further found that CEBPD was inhibited and HMGB3 was highly-expressed in cancer samples of BRCA, and BRCA patients with low expression of CEBPD have poor prognosis (P = 0.025), as well as BRCA patients with high expression of HMGB3 (P = 0.017). (f) Discovery of compound synergy in complex systems intervening in LUAD and LUSC based on SETComp. Based on the predictions of the SETComp model, multiple compounds in Astragalus synergistically inhibit the expression of SPP1 in A549, which was also highly expressed in cancer samples in both LUAD and LUSC, while high expression of SPP1 in LUAD (P = 0.015) and LUSC (P = 0.052) patients has been found to be associated with poor prognosis.

### NP repositioning

Next, we explored the potential of SETComp in the field of drug repositioning. We analyzed several NPs with substantial clinical reports, including ginseng, American ginseng, red ginseng, and cinnamon, and, after masking the clinical reports in advance, we observed whether the model could predict repositioning with clinical evidence. We first performed Kyoto Encyclopedia of Genes and Genomes (KEGG) pathway analysis and Disease Ontology (DO) analysis on the model’s output. Based on different KEGG disease classifications, we organized the potential diseases that these NPs may intervene in (Fig S22a-d). According to the predicted KEGG and DO disease terms, red ginseng was found to liver function-related issues like hepatitis (DOID:1575, hsa05160, has05161) validated by clinical trial NCT0395412, and certain glucose-related diseases like Type II diabetes mellitus (hsa04930), Insulin resistance (hsa04931) and hyperglycemia (DOID:4195), validated by many trials like NCT03775733 and NCT01911663, as well as blood pressure-related diseases like essential hypertension (Fig 5c). American ginseng was found to potentially affect hypertension, multiple sclerosis, respiratory infection and upper respiratory tract infections, which were also found in many clinical trials (Fig 5d), while cinnamon potentially intervened diseases related to diabetes or insulin resistance (Fig S22e) and ginseng potentially affected various diseases, including Alzheimer’s disease, atherosclerosis, glucose-related disorders, liver function issues, and multiple sclerosis (Fig S22f).

### Discovery of compound synergy in NP

Finally, for the most critical issues of complex systems, we provide a potentially feasible application based on our model. We attempted to establish the relationship between the NP-cell-gene associations predicted by the model and the compound-cell-gene associations of the compounds that make up the NP (Fig S23a, b). In the prediction results of Astragalus intervention in MCF-7, we found that the compounds CID: 46906036 and CID: 644263 potentially exert synergistic effects on the gene CEBPD by jointly promoting the expression of CEBPD (Fig 5e), contributing to the upregulation of CEBPD expression in MCF-7 induced by Astragalus. Furthermore, we observed that CEBPD is significantly down-regulated in cancer tissues corresponding to Breast Cancer (BRCA) in MCF-7, and its low expression predicts a significantly worse prognosis (HR=0.69, p=0.025). Similarly, the compounds CID: 3083834 and CID: 644263 in Codonopsis, potentially synergistically inhibit HMGB3 (Fig 5f), which is highly expressed in tumor tissues of BRCA, and patients with high expression of HMGB3 have a significantly poorer prognosis (HR=1.5, p=0.017). Besides, in the prediction of Astragalus intervention in MCF-7, four compounds including CID: 644263 synergistically regulate TPD52 (Fig S23c), a gene that is highly expressed in BRCA tumor tissues, and its high expression is associated with poor prognosis in patients. This suggests that genes such as CEBPD and TPD52 may be potential targets for Astragalus to improve the prognosis of BRCA patients, while the compound CID: 644263 could be a key compound in Astragalus for combating BRCA. On the other hand, in the prediction of Astragalus intervention in A549, compounds CID: 73611 and CID: 5318203 potentially synergistically inhibit the expression of the gene SPP1 (Fig 5g), which is significantly upregulated in both Lung Adenocarcinoma (LUAD) and Lung Squamous Cell Carcinoma (LUSC), the two types of non-small cell lung cancer corresponding to A549. Additionally, in patients with LUAD and LUSC, high expression of SPP1 is associated with poorer prognosis, enhancing its potential as a target for intervention in LUAD and LUSC. The downregulation of the SPP1 gene was also observed in the transcriptomic analysis of Astragalus intervention in A549 (P=0.04, log2FoldChange=-0.28). Furthermore, there are some potentially more complex compound synergies. For example, in the prediction of Astragalus intervention in A549, compounds such as CID: 442813 potentially regulate B3GNT3 (Fig S23d), a gene that is highly expressed in LUSC and associated with poor prognosis in patients with high expression.

## Discussion

As the demand for precision medicine continues to increase^40–44^, the intervention of a single drug has gradually become insufficient to achieve perfect precision medicine, while complex systems composed of multiple compounds offer a promising and potential direction for precision medicine^45^. However, despite the accumulation of vast amounts of multi-omics data from single-compound interventions^46^ and the development of numerous deep models focused on single-compound predictions and generation^47–49^, data reflecting the effects of complex system interventions and models capable of predicting or generating complex system-related functions are scarce. Against this backdrop, there is an urgent need to develop a model capable of predicting the intervention effects of various complex systems, such as NP, on different cell conditions.

Thus, we proposed a model named SETComp, which is capable of predicting genome-wide, cell-specific directed intervention effects for complex systems, such as NP. The two versions of the model (the Concat version with ~200M parameters and the Add version with ~173M parameters) are based on transfer learning and permutation-invariance, with pre-trained compounds as single-element set representations from 970,481,750 compound-cell-gene associations, and further fine-tuned on 2,579,488 compound-cell-gene associations derived from GEO data and literature. The model achieved an accuracy of 93.86% and 92.70%, AUC of 0.9888 and 0.9856, respectively, on the test set of complex system-cell-gene associations. In the ablation experiments, we demonstrated the improvement in model prediction performance brought by permutation-invariance, and in tasks involving complex systems that the prediction model had never encountered before, we achieved prediction accuracies of 82.75% and 82.66%. We also explored the relationship between the model’s predicted output and the true fold change, and found a strong correlation between them. This provides a new perspective for the subsequent development of quantitative prediction or generation models. The model was further validated through in vitro real-world transcriptomic assays that we personally conducted, achieving an accuracy of up to 88.65% among the intervention of multiple NP. We further demonstrated SETComp’s potential in various biomedical application scenarios, including mechanism uncovering of NP, repositioning of NP, and discovery of compound synergistic effects.

There were still some limitations in our studies, including the lack of consideration for the quality ratio of each compound in the complex system, and the inability to achieve quantitative prediction of complex system-cell-gene associations. Fortunately, the set embedding module based on set-based composition, compared to directly summing the feature coefficients of the compounds that make up the complex system (which assumes each compound has the same proportion), can, to some extent, address this issue through various attention mechanisms. Regarding quantitative prediction, although we have not directly achieved quantitative prediction of complex system-cell-gene associations, we have deeply explored the relationship between the model’s predicted output and the actual fold change, observing a strong positive correlation. This provides a promising perspective for the future development of quantitative prediction or generation models.

For future research, on one hand, we plan to collect more single-cell level compound intervention effect data based on the recently published Tahoe-100M dataset^46^, combined with the current 180M-level cell-line-based compound intervention expression profiles, for pre-training generative models. On the other hand, we are currently conducting single-cell sequencing experiments on the intervention effects of multiple complex systems in animals to provide data from a single-cell perspective, which will be used to fine-tune the generative model, ultimately achieving the generation of cell-specific whole-genome expression profiles after complex system interventions. We believe that such a generative model for cell-specific whole-genome expression profiles of complex system intervention will provide better basis for prediction in multiple biomedical application scenarios, such as complex system mechanism uncovering, repositioning, and compound synergy discovery.

## Methods and materials

### Construction and training of the SETComp model

#### Data collection, preprocess and generation

In this work, we collected transcriptomics data of various compounds on different cell lines in CMap from LINCS in NIH Common Fund program^15^, preprocessing with python package cmapPy v4.0.1, and the data was then enhanced using a modified CycleGAN model^18^ to improve the feature dimension from the original 978 to the genome-wide 23,614. Consider the mapping function *G* : *X* → *Y*, where *X* represents the original L1000 assay and *Y* denotes the inferred RNA-seq assay, along with its corresponding discriminator *D*_*Y*_. Accordingly, the adversarial loss function is defined as:

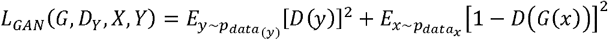

Finally, 1,805,898 samples with a single intervention of 39,321 compounds were used for further study. Compounds with at least 3 technical replications in one cell-line were kept. And the compounds in the processed matrix were annotated in PubChem database^20^ and the chemical structures of these annotated compounds were obtained, represented by SMILES. NP were collected in HERB (v2.0)^21^ database, and information like chemical compositions and Latin names were collected. Compounds in these NP were also annotated in PubChem database for standard PubChem CID and chemical structures. A total of 46,419 compounds remained, among which 25,751 served as chemical compositions in at least one of 6198 NP.

For compounds, the enhanced transcriptomic data uniformly after 24h of intervention at the highest concentration of the compound was uniformly used to calculate expression pattern of each gene, while transcriptomic data or microarray in which cell lines were intervened by NP were collected from GEO database (update to 2024 May) and literature^50,51^. The expression data was further processed with limma^52^ and DESeq2^53^ to calculate DEGs with statistical models. The expression pattern of each gene in different cell lines under different interventions by compounds or NP was classified into 3 categories, including Up (log2FoldChange > 0, *P* value < 0.05), Down (log2FoldChange < 0, *P* value < 0.05) and No effect (*P* value > 0.05).

The data and information of cell lines were collected from CCLE and TCGA database. The expressions matrix of CCLE and TCGA were processed with log transform, if necessary. The data and information of genes were collected from STRING database^19^, including the interactions between two genes and the corresponding encoded protein sequence. Finally, 970,481,750 compound-cell line-gene pairs and 2,579,488 NP-cell line-gene pairs were obtained for further data pre-training and fine-tuning in 3 classes (up-regulated, down-regulated and no change), respectively.

#### Feature extraction for compounds, genes and cell lines

Based on the processed data, every compound was firstly embedded in a self-supervised pre-training model, Infograph^54^, which proposed to maximize the mutual information between the graph-level and node-level representations and was implemented in TorchDrug. In the InfoGraph model, the mutual information estimator *I* _*ϕ*_,_*ψ*_, is modeled using the discriminator, *T* _*ψ*_ which is parameterized by a neural network with parameters *ψ*:

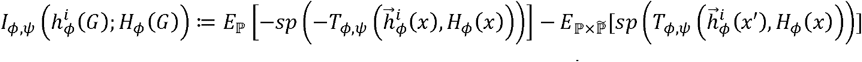

Here, *x* denotes an input sample, while the negative sample *x*^′^ is drawn from the distribution 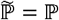, which matches the empirical distribution of the input space. Additionally, the softplus function is defined as *sp*(*z*) = log (1 + *e*^*z*^). And the InfoGraph model was trained on the ZINC 2M database using the recommended parameters, with five hidden layers and 300 neurons in each layer. Another embedding for compounds was based on the PubChem fingerprint calculation implemented with scyjava package in Python. Finally, each compound was represented by a 1,181-dimensional feature vector, consisting of an 881-dimensional fingerprint embedding and a 300-dimensional Infograph embedding. The features of NP were composed of the corresponding features of chemical compositions and assembled into a set for each natural product.

Genes were also embedded in two ways, including Node2Vec based on network relationships and ProtFlash^55^, A lightweight protein language model, based on corresponding encoded protein sequence. The loss function of the masked training of ProtFlash is defined as:

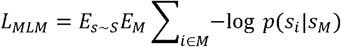

In which *s*_*i*_ is the true amino acid and *s*_*M*_ is the masked sequence as context. Node2Vec algorithm was implemented by torch geometric to achieve parallel training on GPU and every gene was embedded into 256 dimensions, while ProtFlash model was used with pre-trained weights and every gene was embedded into 768 dimensions.

For cell line embedding, a Variational Autoencoder (VAE) model was first trained on the TCGA expression matrix, and the CCLE expression matrix was then embedded with the trained VAE model. The loss of the VAE model is defined as the sum of MSE loss and KL loss:

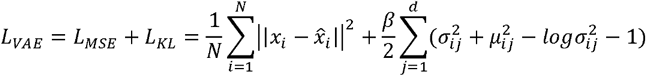

In this function, *N* represents the total number of samples in the dataset, where each sample *x*_*i*_ is the original expression profile and 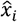 is the corresponding reconstructed profile generated by the decoder. For each sample, the approximate posterior distribution is characterized by the mean *μ*_*ij*_ and standard deviation *σ*_*ij*_ for the *j*-th dimension of the latent variable *z*, with *d* denoting the overall dimensionality of the latent space. A hyperparameter *β* is introduced to control the weight of the KL divergence term, allowing for a balance between reconstruction fidelity and latent space regularization.

Grid search was applied to find the best hyper-parameter combination, in which the search ranges were set as follows: batch size: 32, 64, 128, 256; learning rate: 1e-3, 1e-4, 1e-5; dimension in MLP layer 1: 4096, 2048, 1024; dimension in MLP layer 2: 1024, 512, 256; dimension in MLP layer 3: 256, 128, 64; dimension in latent layer: 64, 32, 16. For each combination, the model was trained for 20 epochs. And the loss function was composed of two parts: Mean Squared Error loss as the reconstruction loss and Kullback-Leibler Divergence loss. Finally, every cell line was embedded into 64 dimensions, in which VAE showed the best reconstruction performance.

#### Main architecture of the model

Apart from the feature extraction modules for compounds, genes and cell lines, the main architecture of the model was constructed with three modules including the set embedding module, the attention module and the prediction module. In the set embedding module, we utilized Deep Sets^16^ and Set Transformer^17^ (See Supplementary materials) to effectively model set-structured data inherent in our problem domain. The set embedding module consisted of two Deep Sets models and one Set transformer model, both implemented in Pytorch environment (https://github.com/juho-lee/set_transformer/tree/master). Traditional neural network architectures often assume a fixed-size input or a specific ordering of elements, which is unsuitable for sets that are inherently unordered and variable in size. Deep Sets address this challenge by providing a framework that is permutation-invariant to the input set elements. For the Concat version and Add version of the SETComp model, the outputs of the set embedding are as follows, respectively:

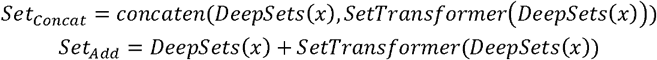

In addition, the attention module further refines the representations obtained from the concatenate of the set embedding module for compounds or NP and the extracted gene embedding and cell line embedding. It utilizes advanced attention mechanisms to focus on the most relevant features within the data, enhancing the model’s predictive capabilities by capturing intricate patterns and dependencies. Finally, the prediction module takes the refined representations from the attention module to perform the final prediction task. This module typically consists of three fully connected layers that map the high-dimensional embeddings to the desired output space, enabling the model to generate accurate predictions based on the learned representations.

### In vitro cell lines intervention by multiple NP for validation

#### Cell Culture and Related Reagents

The human-derived breast cancer cell line MCF-7 and human-derived lung cancer cell line A549 were both purchased from the China Infrastructure of Cell Line Resources (China). These cells were then cultured in DMEM and RPMI-1640 media, respectively, with 10% (v/v) FBS and 100 U/ml streptomycin/penicillin at 37°C in a 5% CO2 environment. The Chinese medicinal herbs Astragalus, Cinnamon, and Codonopsis were purchased from Jiangyin Tianjiang Pharmaceutical Co., Ltd. (China). All the herbal granules were stored in a dry, light-protected environment.

#### Cell Treatment and Drug Administration Method

Human-derived breast cancer cells (MCF-7) and human-derived lung cancer cells (A549) were digested with trypsin, resuspended, and 10 μL of cell suspension (1:1 dilution) was counted using a cell counting plate. The cells were then seeded in a 10 cm cell culture dish at a density of 4 × 10^6 cells per well, and placed in the cell incubator to allow attachment. The following day, after the cells attached, the culture medium was replaced with 8 mL of fresh serum-free medium. Astragalus, Cinnamon, and Codonopsis were adjusted to final concentrations of 0, 100, and 200 μ g/mL, respectively, and then cultured in a 37°C, 5% CO2 cell incubator for 24 hours. On the third day, cells from each group were collected into 1.5 mL sterile, enzyme-free, Eppendorf tubes using Trizol reagent, and stored at −80°C for sequencing.

#### RNA extraction, library construction, and sequencing

Total RNA was extracted from the cell lines using TRIzol® reagent (Magen). The A260/A280 absorbance ratio of the RNA samples was measured using a Nanodrop ND-2000 (Thermo Scientific, USA), and the RNA Integrity Number (RIN) was determined using an Agilent Bioanalyzer 4150 (Agilent Technologies, CA, USA). Only RNA samples that passed quality control were used for library construction. The PE library was prepared according to the instructions of the ABclonal mRNA-seq Lib Prep Kit (ABclonal, China). mRNA was purified from 1μg of total RNA using oligo(dT) magnetic beads, and then fragmented in the ABclonal First Strand Synthesis Reaction Buffer. Subsequently, mRNA fragments were used as templates to synthesize the first strand of cDNA using random primers and reverse transcriptase (RNase H). The second strand of cDNA was synthesized using DNA polymerase I, RNase H, buffers, and dNTPs. The synthesized double-strand cDNA fragments were ligated with adapter sequences for PCR amplification. The PCR products were purified and the library quality was assessed using an Agilent Bioanalyzer 4150. Finally, sequencing was performed on the Illumina Novaseq 6000 / MGISEQ-T7 sequencing platforms.

### Application of the SETComp model in biomedical scenarios

#### Mechanism uncovering for NP

For the prediction of complex system intervention effects, we selected genes from class 0 and 1 with softmax scores greater than or equal to 0, 0.7, and 0.9 as potential upregulated or downregulated genes under different thresholds. These genes were then sorted by their scores (with negative scores assigned to genes predicted as class 1) for subsequent enrichment analysis. Next, we performed GSEA analysis using the R package clusterProfiler (v4.9.1) to identify pathways potentially intervened by the model, and classified the pathways into activation or inhibition based on Normalized Enrichment Score (NES). This enrichment analysis was repeated for potential intervention genes under different thresholds to obtain mechanism uncovering results for each threshold. For visualizing specific pathways, we used the gseaplot2() function from the R package enrichplot.

#### Repositioning for NP with clinical evidences

We collected clinical reports of various complex systems from the HERB (v2.0) database to validate our findings in drug repositioning. Based on the predicted effects of complex system interventions, we selected genes with softmax scores greater than or equal to 0.7 as potential intervention genes. Using the R package clusterProfiler (v4.9.1), we performed KEGG enrichment analysis and DO enrichment analysis on the potential intervention genes to identify diseases or disease-related terms potentially intervened by each complex system. These disease or disease-related terms were then modularized according to KEGG classifications. We matched the disease or disease-related terms with clinical diseases and observed whether there were relevant clinical trial reports. Sankey diagrams were visualized using the R package ggalluvial (v0.12.5).

#### Gene-level compound synergistic effects finding in NP

For the prediction of compound intervention effects and complex system intervention effects, we selected genes from class 0 and 1 with softmax scores greater than or equal to 0.7 as potential upregulated or downregulated genes. Tumor and normal samples of BRCA, LUAD, and LUSC were obtained from the UCSC Xena project^56^, which has normalized the expression matrices of TCGA tumor samples and GTEx normal samples. After processing with DESeq2, differential genes for BRCA, LUAD, and LUSC were obtained. Based on the clinical prognostic information of individual tumor patients from TCGA and the GEPIA2 platform^57^, we analyzed the prognosis differences between high and low expression groups of each gene (grouped by the median gene expression).

## Supporting information

Supplementary meterials

## Code availability

The code related to this manuscript has been uploaded to GitHub and set to private. It will be made public after the manuscript is officially submitted for review.

## Acknowledgements

This work was supported by National Administration of Traditional Chinese Medicine (GZY-KJS-2024-03).

## Author contributions

S.L. contributed to the conception and design of the work. B.W. developed the concept of the work, contributed to design and implementation of the algorithm and drafted the initial version of the manuscript. P.Y. contributed to conducting experimental validation. B.W., T.Z., and Q.L. contributed to the data collection and preprocessing.

