## Supplementary meterials for "Transfer Learning and Permutation-Invariance improving Predicting Genome-wide, Cell-Specific and Directional Interventions Effects of Complex Systems"

### Methods and materials

#### Construction and training of the SETComp model

##### Training details for pre-training and fine-tuning of the model

Our goal is to predict the up- or down-regulatory effect of given compounds or NP on every gene of a certain cell line. So, the model is trained based on a common classification loss with L2 regulation. Given the total number of training samples $N$, and the number of classes in the classification task $K = 3$, for each sample, $y_{ij}$ is the ground-truth label for class $j$ and $\hat{y}_{ij}$ is the predicted probability for class $j$:

$$L = -\frac{1}{N}\sum_{i=1}^{N}\sum_{j=1}^{K} y_{ij} log\hat{y}_{ij} + \lambda||\theta|\ |_2^2$$

In the pre-training procedure, 28,627,074 compound-cell line-gene pairs, in which compound served as chemical compositions in at least one natural product, were extracted as the test dataset. To address the potential issue of imbalance among samples, we applied a down-sampling method, reducing the sample sizes of the other two categories to match the size of the smallest category. And the training and validation sets were divided in a 7:3 ratio. Input features of training data, validation data and test data were scaled on training data for compounds and genes, respectively. Grid search was applied to find the best hyper-parameter combination, in which the search ranges were set as follows: batch size: 1024, 512, 256, 128, 64; learning rate: 1e-3, 1e-4, 1e-5, 1e-6; L2 regularization: 1e-5, 1e-4, 1e-3, 1e-2; dropout: 0.1, 0.2, 0.5, 0.8; dimension in MLP layer: 512, 1024; dimension in Set Transformer layer: 512, 1024, 2048. For each combination, the model was trained for 5 epochs. And for the add version, we used the same structure for grid search, while L2 regularization and dropout were set as the varied parameters: L2 regularization: 1e-5, 1e-4, 1e-3, 1e-2; dropout: 0.1, 0.2, 0.5, 0.8.

In the fine-tuning step, the down-sampling method was also performed, while the training, validation and test sets were divided in a 7:2:1 ratio. Input features of training data, validation data and test data were scaled on training data for compounds and genes, respectively. Another grid search was performed, in which the search ranges were set as follows: batch size: 1024, 512, 256, 128, 64; learning rate: 1e-3, 1e-4, 1e-5, 1e-6; L2 regularization: 1e-5, 1e-4, 1e-3, 1e-2; dropout: 0.1, 0.2, 0.5, 0.8. For each combination, the model was trained for 50 epochs with an early stopping strategy, setting patience as 5 epochs.

The model was trained in two 4090 GPUs with DataParallel implemented in PyTorch.

##### Main architecture of the model

The core idea of Deep Sets is to encode a set $X$ by applying a shared transformation $\varphi$ to each element and then aggregating the transformed elements using a permutation-invariant operation $\rho$ such as summation or averaging. Mathematically, this can be expressed as:

$$f(X) = \rho(\{\varphi(x_1), \varphi(x_2), ..., \varphi(x_n)\})$$
$$X = \{x_1, x_2, ..., x_n\}$$

This formulation ensures that the output representation $f(X)$ remains unchanged under any permutation of the input set elements, capturing the essence of the set's properties without imposing any arbitrary ordering.

Building upon the foundation of Deep Sets, Set Transformer introduce a more expressive and scalable approach by incorporating self-attention mechanisms, inspired by the success of transformers in sequence modeling. Set Transformer are designed to handle sets by using attention

layers that are permutation-invariant and can model complex interactions between set elements.
The Multihead Attention Block (MAB) is the core component of the Set Transformer. MAB combines queries $Q$, keys $K$, and values $V$, leveraging the multi-head self-attention mechanism to capture relationships between elements. It is defined as:

$$MAB(Q,K,V) = LN(Q + MultiHead(Q,K,V))$$

where $LN$ denotes Layer Normalization, and $MultiHead$ represents the multi-head attention mechanism, computed as:

$$MultiHead(Q,K,V) = Concat(head_1, ..., head_h)W^O$$

Each attention head $head_i$ is calculated as:

$$head_i = Attention(QW_i^Q, KW_i^K, VW_i^V)$$

The Set Attention Block (SAB) is composed of MABs and is used to perform self-attention operations on the input set. SAB is formulated as:

$$SAB(X) = MAB(X,X,X)$$

where $X$ represents the input set's representation. Through SAB, the model can capture higher-order interactions among set elements.
The Induced Set Attention Block (ISAB) introduces the concept of inducing points to reduce computational complexity. ISAB uses a fixed number of inducing points to map the input set to a lower-dimensional space, thereby reducing computation while maintaining model performance. It is defined as:

$$H = MAB(I,X)$$

$$ISAB(X) = MAB(X,H)$$

where $I$ is the learnable inducing point matrix, and $H$ is the intermediate representation.
Pooling by Multihead Attention (PMA) is used to aggregate variable-length set representations into a fixed-size global representation. PMA employs a set of $k$ learnable seed vectors, interacting with the input set through the multi-head attention mechanism, defined as:

$$PMA_k(X) = MAB(S,X)$$

where $S$ is a set of $k$ learnable seed vectors, and $k$ is the desired number of output elements.
By stacking these attention blocks, Set Transformer captured higher-order interactions among elements without assuming any particular sequence, making them highly suitable for tasks involving set inputs. The self-attention mechanism allows the model to dynamically weigh the relevance of each element relative to others, resulting in a richer representation of the set. In general, we leveraged these architectures to process inputs naturally represented as sets, ensuring that our models remained invariant to the permutation of input elements while effectively capturing complex relationships within the data. This approach enhances the model's ability to generalize and accurately reflect the underlying structure of the data without introducing biases associated with input ordering.

**Baseline model comparison and ablation studies**

In comparison to our proposed model in both predicting the targets of single compounds and NP, we applied baseline models for pre-training and fine-tuning, respectively. For pre-training procedure, we trained classic machine learning models including K-Nearest Neighbors (KNN) and Decision Tree (DT), Linear Discriminant Analysis (LDA), as well as deep learning models for ablation studies, including Vanilla Neural Network (MLP), MLP with attention module (named No-set), and model with same structures but trained on smaller training datasets. KNN and DT were constructed based on python package scikit-learn. As in the pre-training procedure, embeddings of compounds were sets containing only one element, so we treated them as 2D vectors which would be directly combined with the embeddings of genes and cell lines as the input features for No-set, MLP, KNN, and DT. Besides, we also trained models on smaller dataset which had 1/100 training samples of the original training dataset. This was done on the Concat version, Add version of our models, as well as No-set, MLP, KNN, and DT. We only test the performance of KNN and DT on the smaller dataset for their low computing efficiency and large demand of memories. In general, in the pre-training step, we directly compared the full version of the Concat version, Add version of our model with No-set on the full training dataset, as well as smaller version of the Concat version, Add version of our model with No-set, MLP, KNN, and DT on the 1/100 dataset. In details, the training parameters for MLP and No-set chose the best grid search parameters of the Concat version in the pre-training grid search.

In fine-tuning procedure, we compared the performance of the Concat version, Add version of our model with No-set, MLP, KNN, DT, and LDA directly trained on the fine-tuning dataset as well as small pre-training dataset with fine-tuning dataset, MLP and No-set trained on pre-training dataset with fine-tuning dataset. Additionally, we also compared our models with the Concat version, Add version of our models only trained on the pre-training dataset, only trained on the fine-tuning dataset, as well as the models trained on the smaller pre-training dataset and full fine-tuning dataset. Specially, as No-set, MLP, KNN, DT, LDA only accepted 2D dimension feature in this circumstance, the set input was averaged into 2D dimension vector within their training or testing. Further we estimated the performance of our models which were trained on the full pre-training dataset and different percentages of the fine-tuning dataset to find the relationship between the performance and the size of the fine-tuning dataset. In details, the training parameters for MLP and No-set chose the best grid search parameters of the Concat version in the fine-tuning grid search. Finally, we also tried another fine-tuning task, in which we divided the whole fine-tuning datasets according to the intervention of different NP. Among the 143 kinds of NP, transcriptomics intervened by 10% of them (14) were chosen as the test data, and the remaining part was split into the training set and the validation set with the ratio of 7:3. The grid search was also performed, in which the search ranges were set as follows: batch size: 1024, 512, 256, 128, 64; learning rate: 1e-3, 1e-4, 1e-5, 1e-6; L2 regularization: 1e-5, 1e-4, 1e-3, 1e-2; dropout: 0.1, 0.2, 0.5, 0.8.

In the comparison to baseline models and the ablation studies, Area under ROC curve (AUC), Area under Precision-Recall curve (AUPR), Accuracy (Acc) and F1 score were chosen as the metrics for comparison. Acc was calculated according to:

$$Acc = \frac{1}{N}\sum_{i=1}^{N} 1(\hat{y}_i = y_i)$$

F1 score was weighted-average, for every class $i$:

$$F1_i = \frac{2 \cdot Precision_i \cdot Recall_i}{Precision_i + Recall_i}$$

$$F1_{weighted} = \frac{\sum_{i=1}^{K} N_i \cdot F1_i}{N}$$

In which:

$$Precision_i = \frac{TP_i}{TP_i + FP_i}$$

$$Recall_i = \frac{TP_i}{TP_i + FN_i}$$

And $TP_i$ is the number of true-positive for class $i$, $FP_i$ is the number of false-positive for class $i$ and $FN_i$ is the number of false-negative for class $i$.

AUC was also weighted-average on every class:

$$AUC_i = \int_0^1 TPR_i(FPR)dFPR$$

$$AUC_{macro} = \frac{1}{K}\sum_{i=1}^{K} AUC_i$$

So as AUPR:

$$AUPR_i = \int_0^1 Precision_i(Recall)dRecall$$

$$AUPR_{macro} = \frac{1}{K}\sum_{i=1}^{K} AUPR_i$$

#### In vitro cell lines intervention by multiple NP for validation

##### Quality control and alignment

The raw data in FASTQ format were first processed using a Perl script to remove adapter sequences and filter out low-quality reads (where the number of bases with quality scores $\leqslant$ 25 exceeds 60% of the total reads) and reads with more than 5% N (N indicates undetermined base information), resulting in clean reads for subsequent analysis. HISAT2[1] software was used to align the clean reads to the reference genome, generating mapped reads for further analysis.

##### Differential expression analysis for validation

To perform differential gene analysis based on the count matrix, we used the R package DESeq2 (v1.30.0)[2]. Additionally, since some samples had low quality when initially submitted, we re-submitted the samples to ensure their quality, and imported batch correction during the DESeq2 differential gene calculation.

Based on DESeq2, we obtained the foldchanges and P values for each transcript (with ENSEMBL as the id) after the interventions of different NP. The ENSEMBL id for each transcript was converted to ENTREZ id and gene symbol by function bitr() in R package clusterProfiler (v3.18.0)[3]. We calculated the accuracy of the model's predictions based on the following formula:

$$Acc_{class_i} = \frac{\left|intersection\left(predict_{class_i}, truth_{class_i}\right)\right|}{\left|predict_{class_i}\right|}, i \in \{0,1\}$$

Here, $predict_{class_i}$ represented the genes predicted and classified by SETComp into $class_i$ with the softmax of the output no less than the defined threshold and $truth_{class_i}$ refers to all genes or differentially expressed genes belonging to $class_i$ determined by foldchanges. We performed the statistical analysis separately for both scenarios under the conditions of each NP intervention in each cell line.

Additionally, for the predictions classified as class 2, which indicates no significant intervention effect, we performed statistical analysis using the following formula:

$$Acc_{class_2} = \frac{\left|intersection\left(predict_{class_2}, ns\right)\right|}{\left|predict_{class_2}\right|}$$

In which, $ns$ represented all the genes with P value no less than 0.05 and $predict_{class_2}$ has two calculation methods: one is for genes predicted to belong to $class_2$, and the other is for genes predicted not to belong to $class_0$ or $class_1$.

**Supplementary Figures**

Supplementary Figure S1

**Supplement Figure S1**

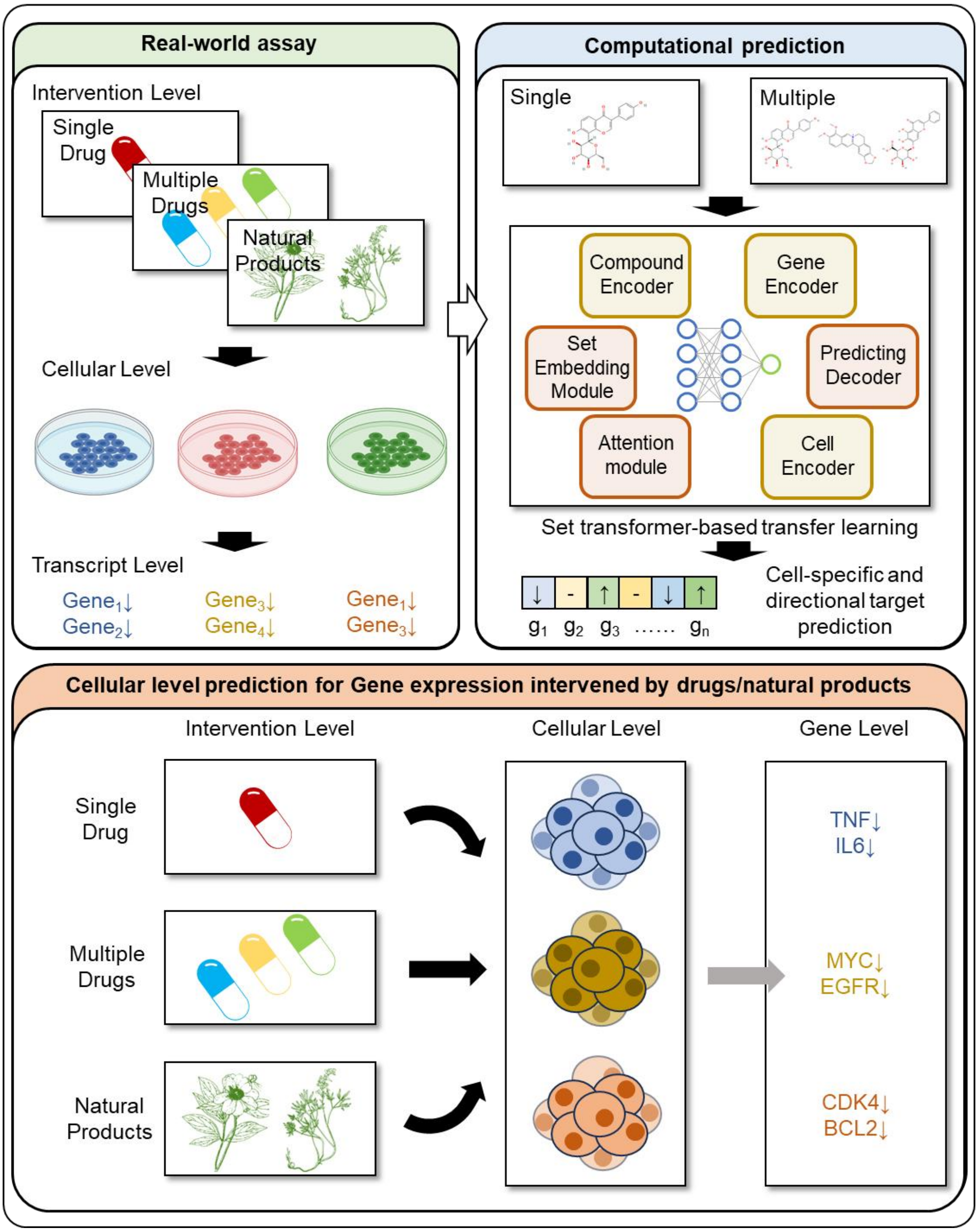

Supplementary Figure S2

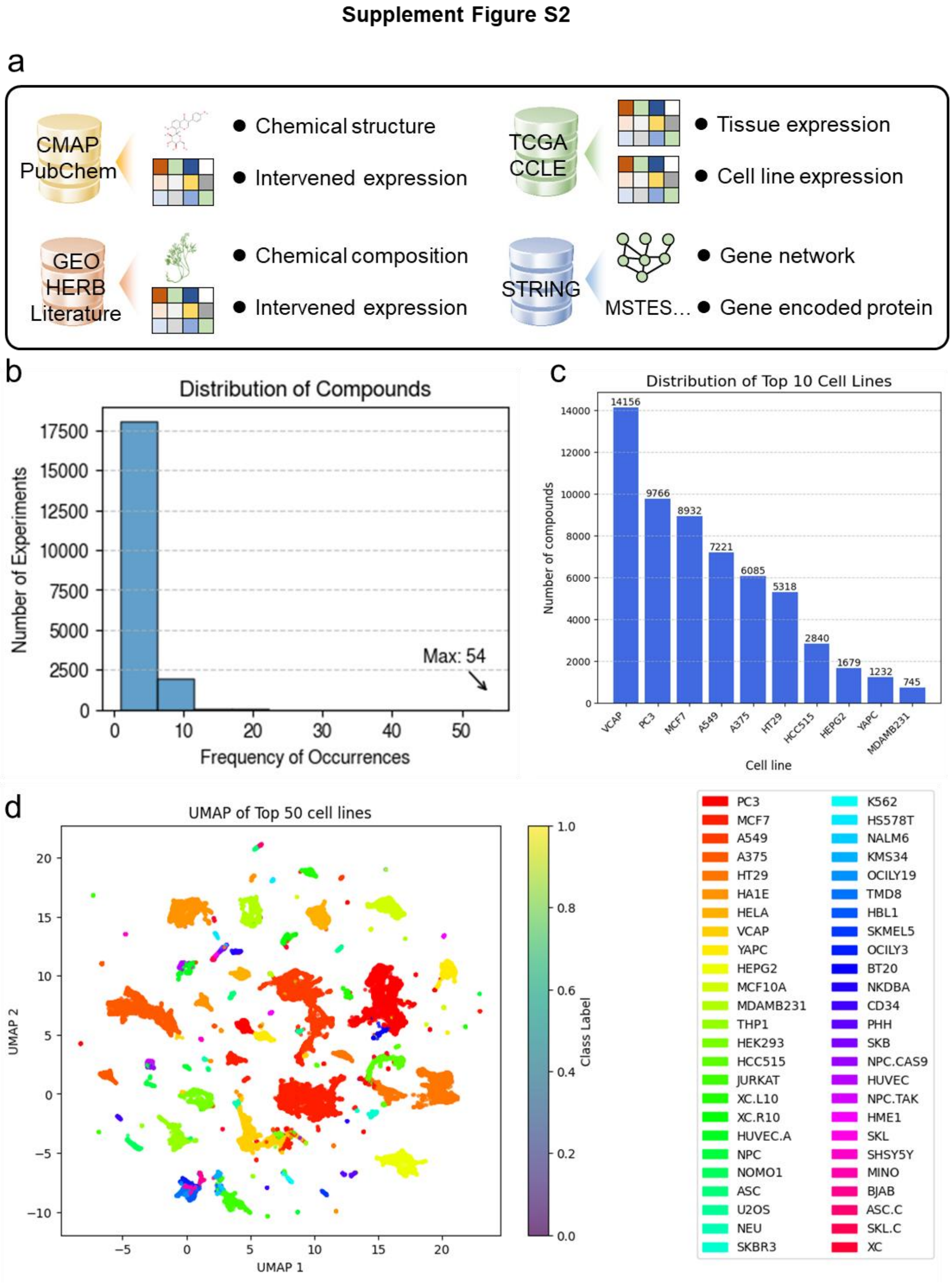

Supplementary Figure S3

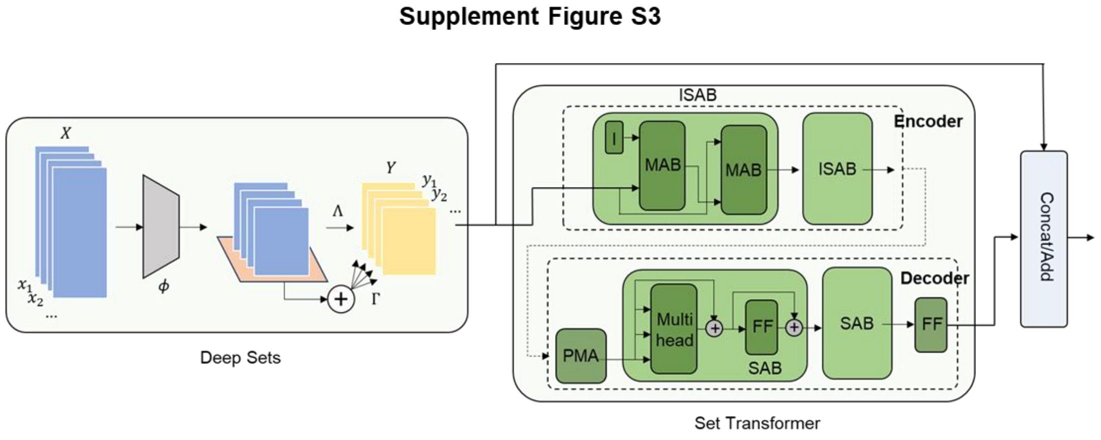

Supplementary Figure S4

**Supplement Figure S4**

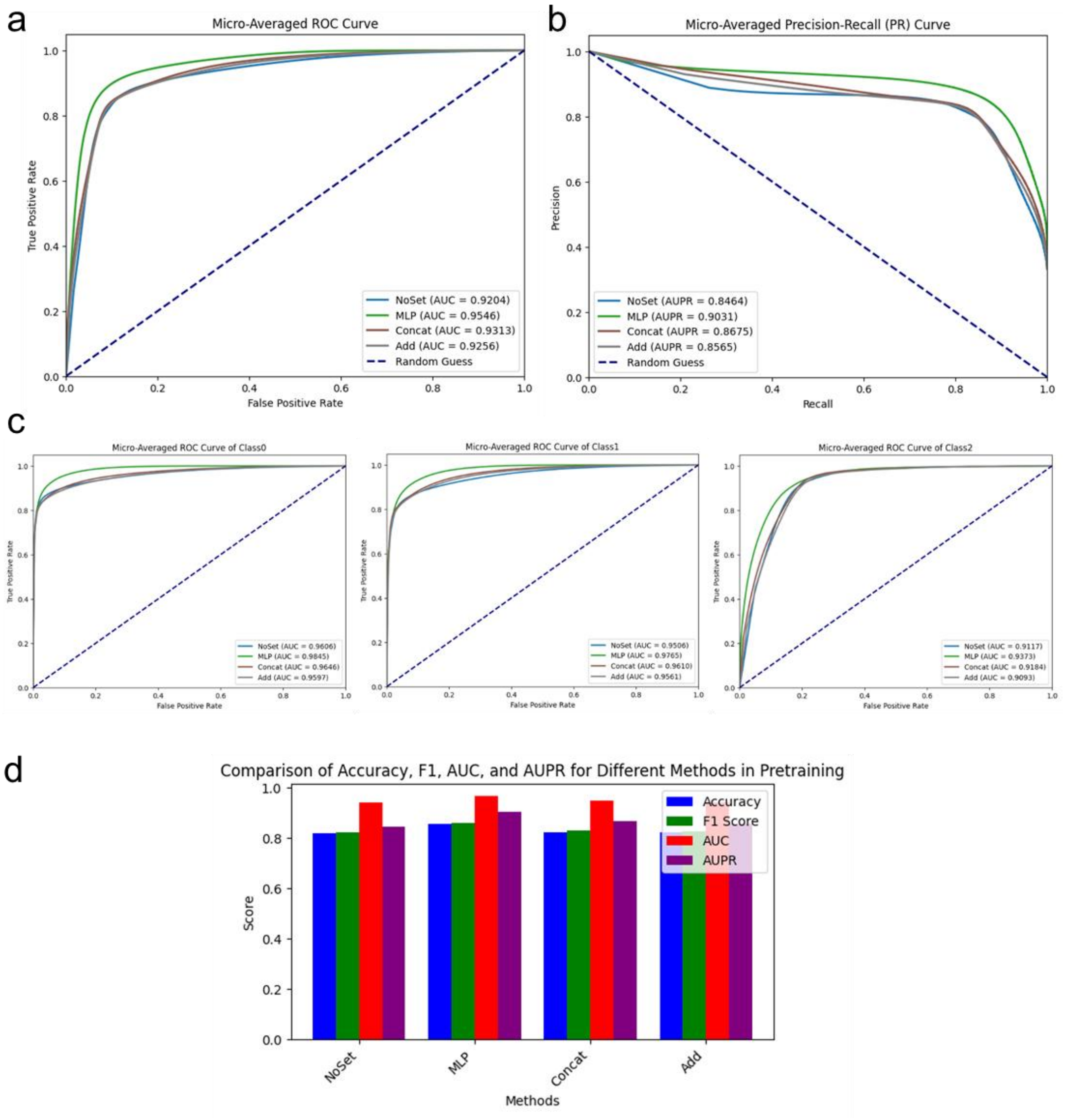

Supplementary Figure S5

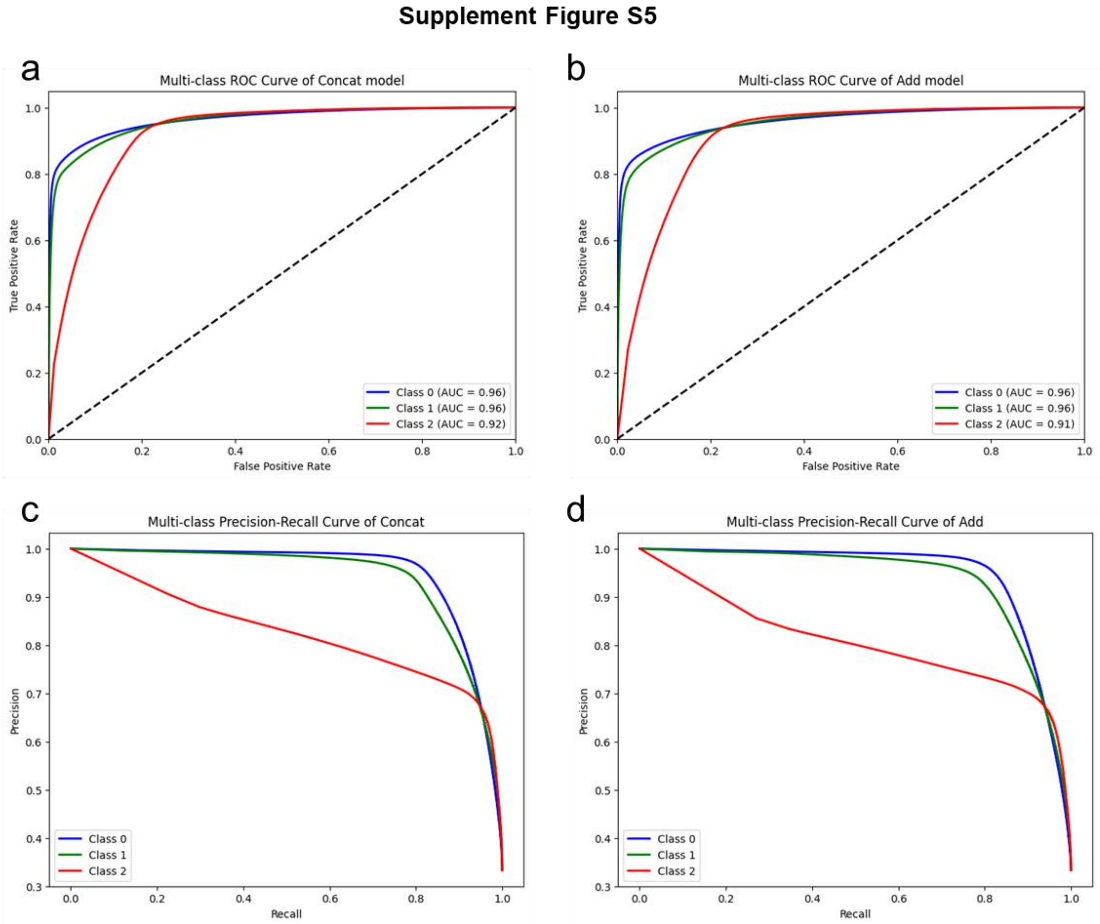

Supplementary Figure S6

**Supplement Figure S6**

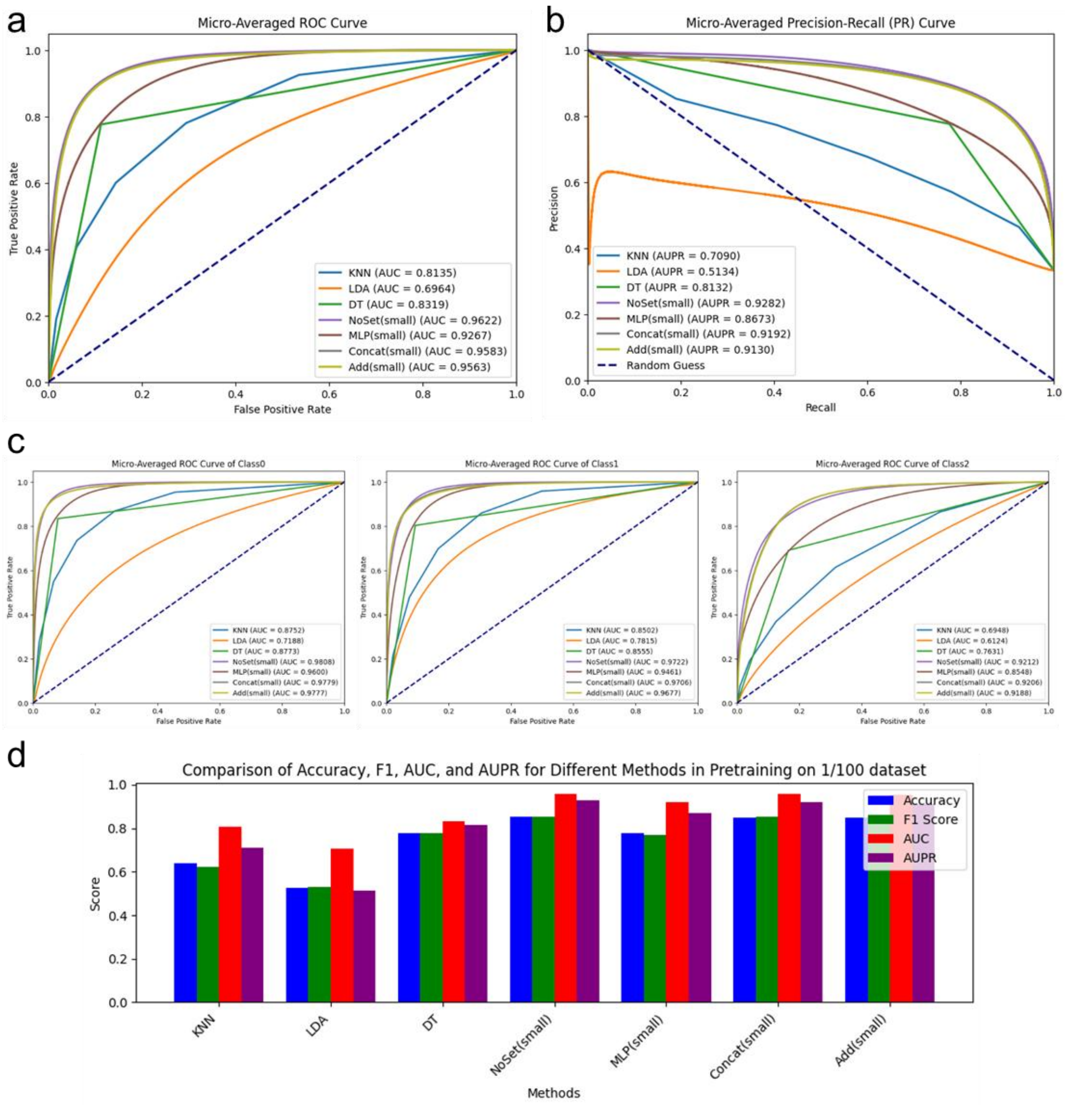

Supplementary Figure S7

**Supplement Figure S7**

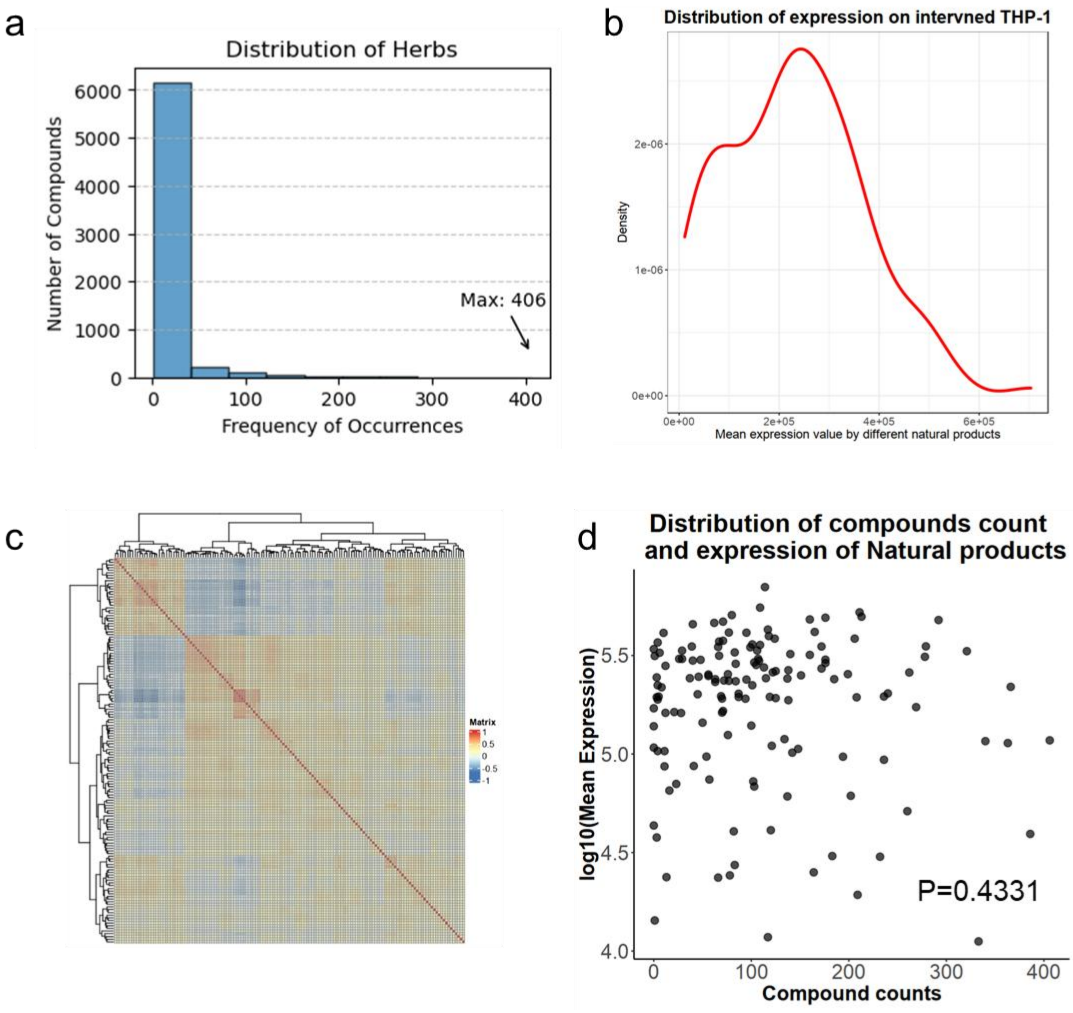

Supplementary Figure S8

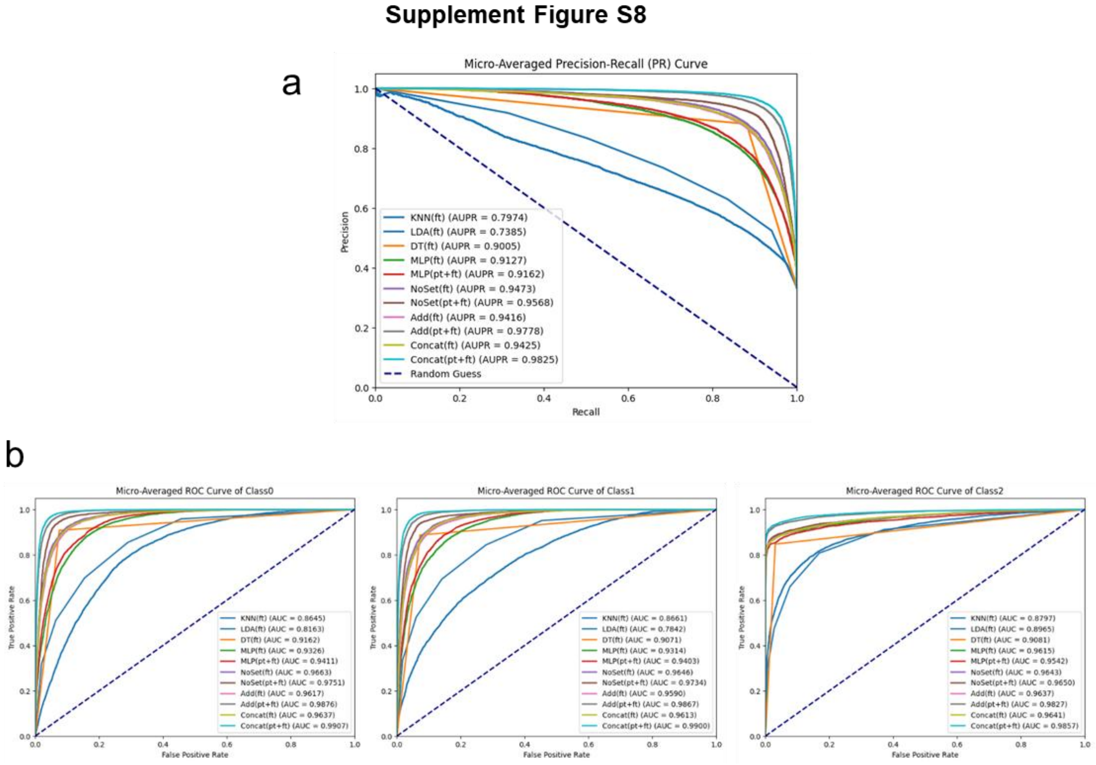

Supplementary Figure S9

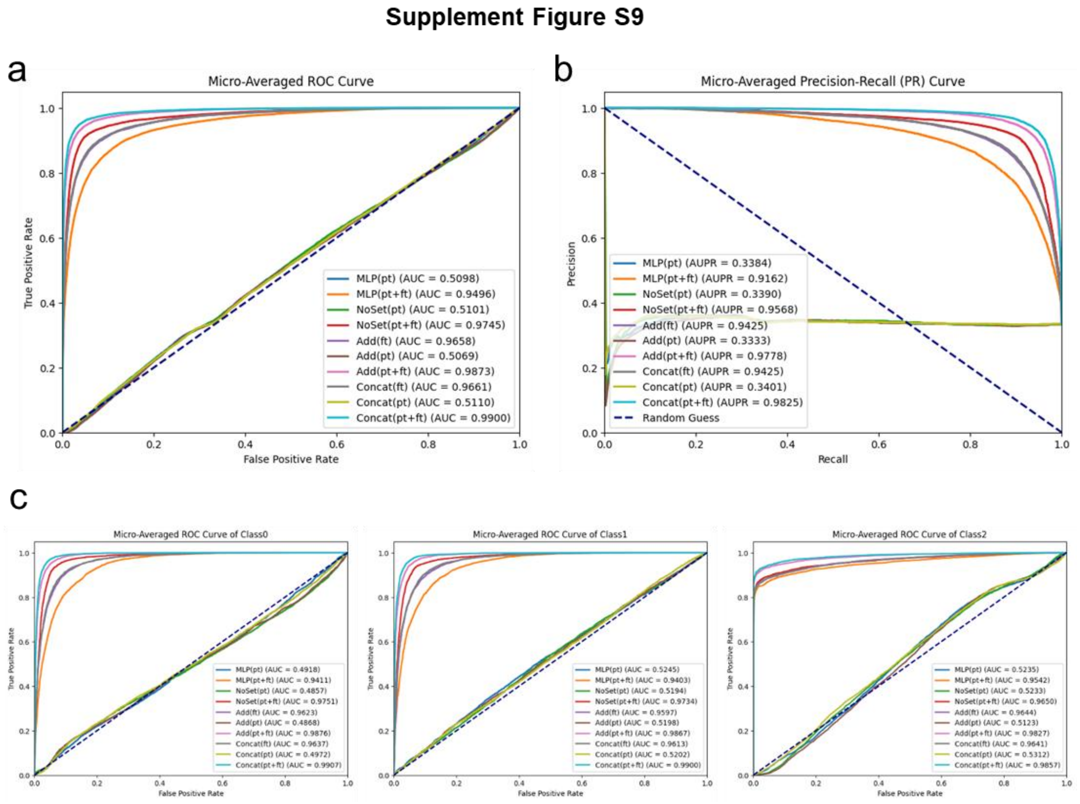

Supplementary Figure S10

**Supplement Figure S10**

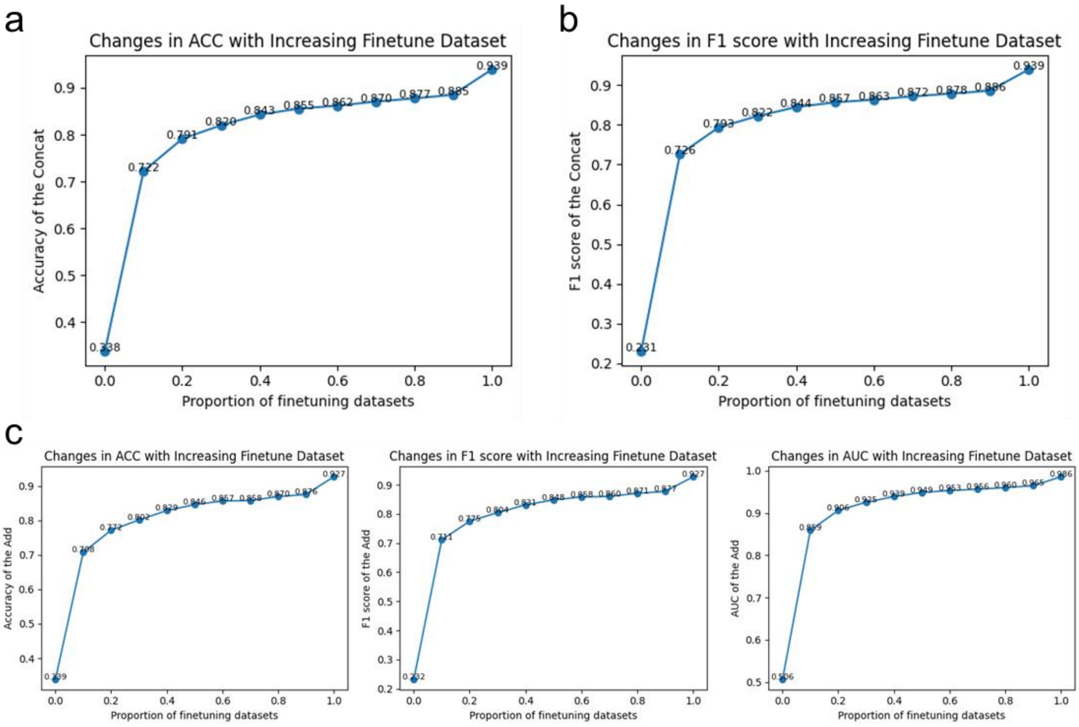

Supplementary Figure S11

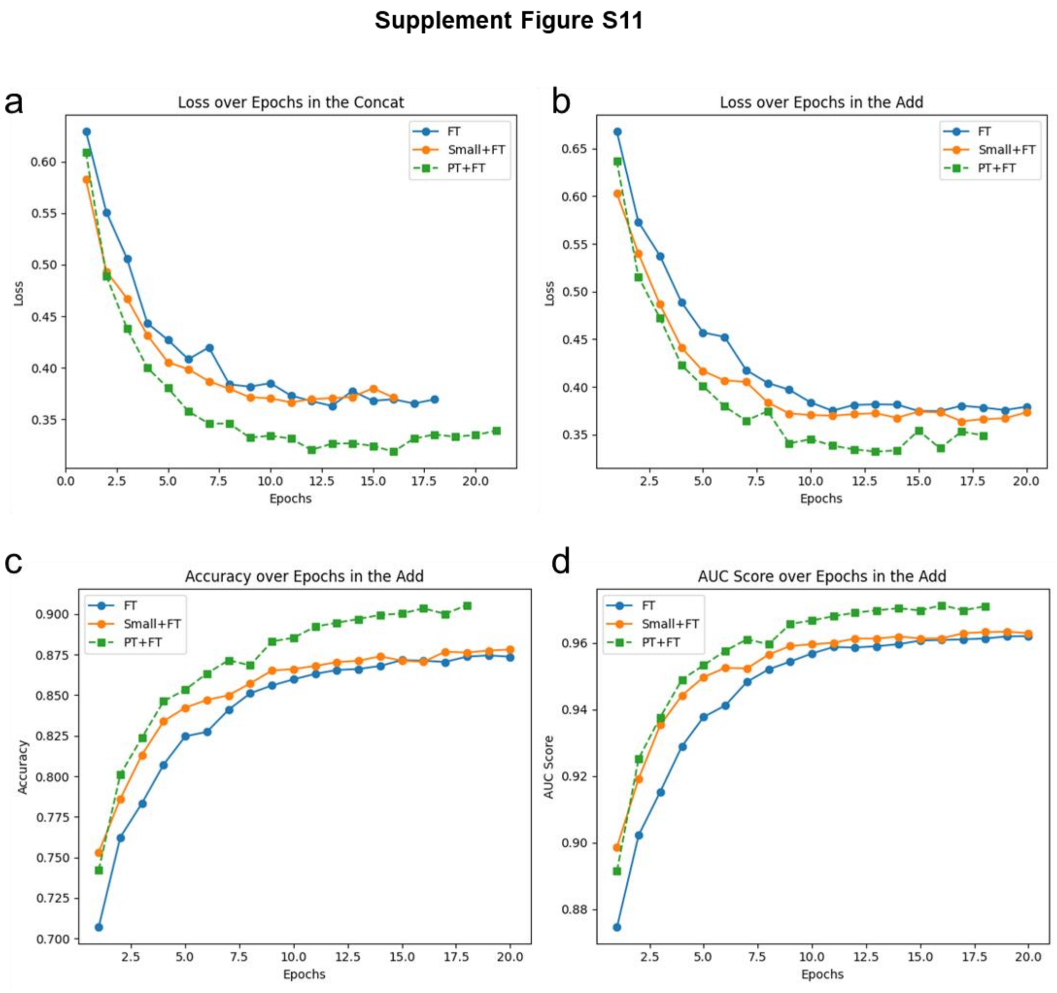

Supplementary Figure S12

**Supplement Figure S12**

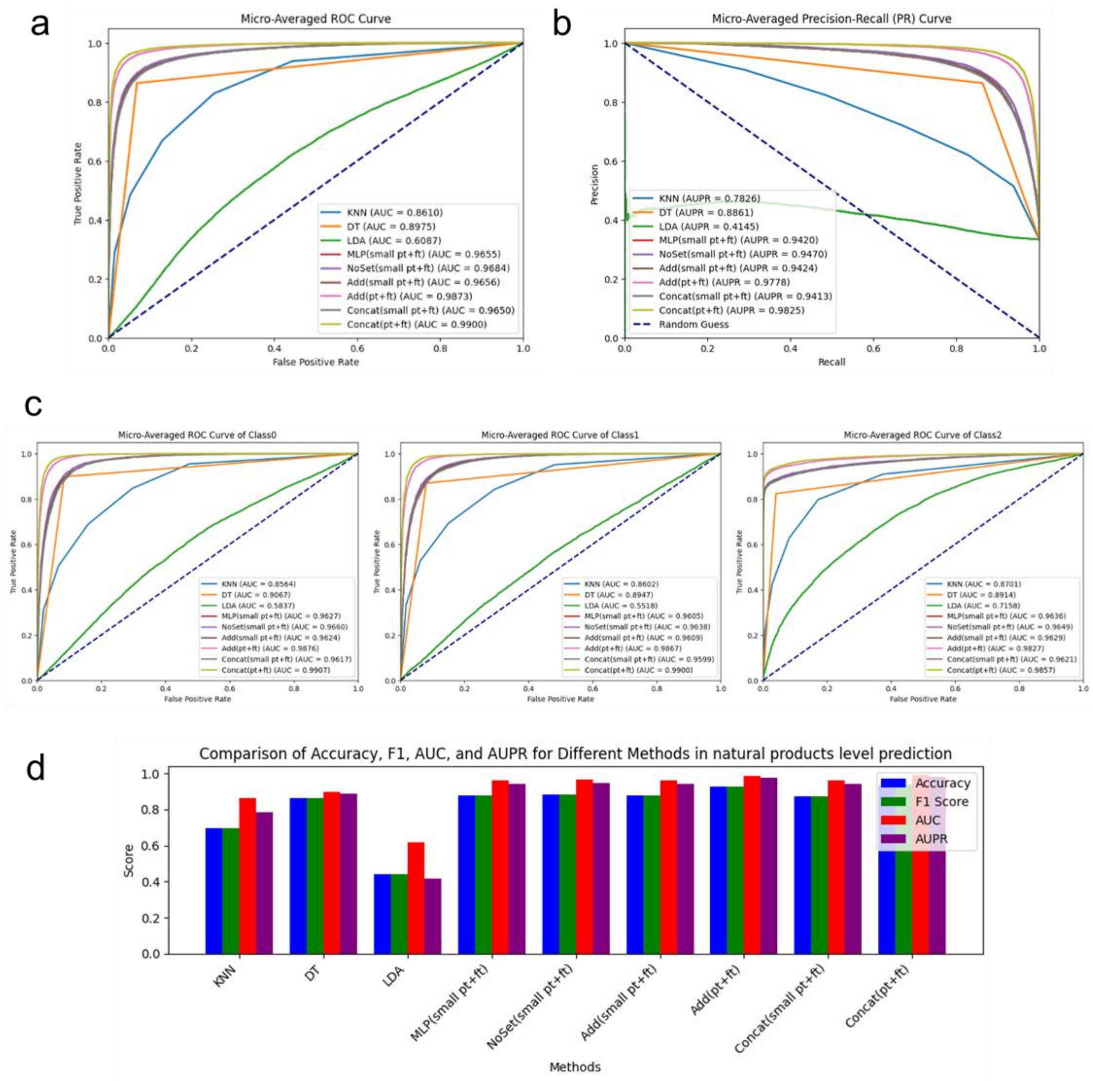

Supplementary Figure S13

**Supplement Figure S13**

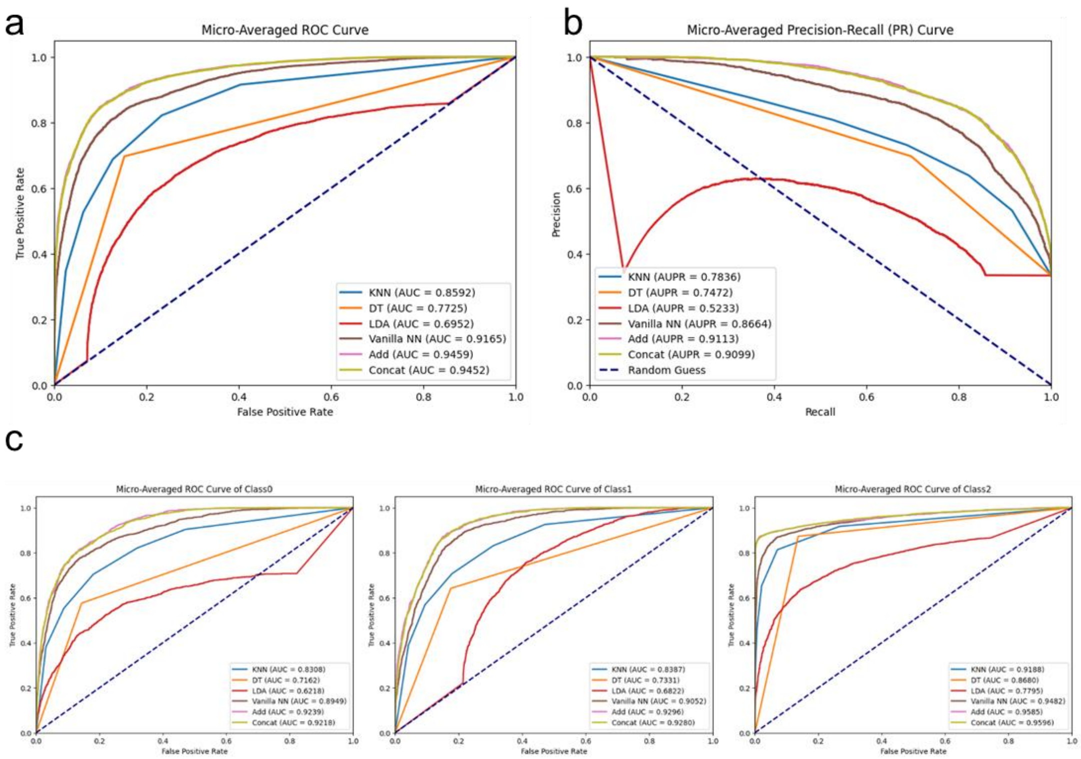

Supplementary Figure S14

#### Supplement Figure S14

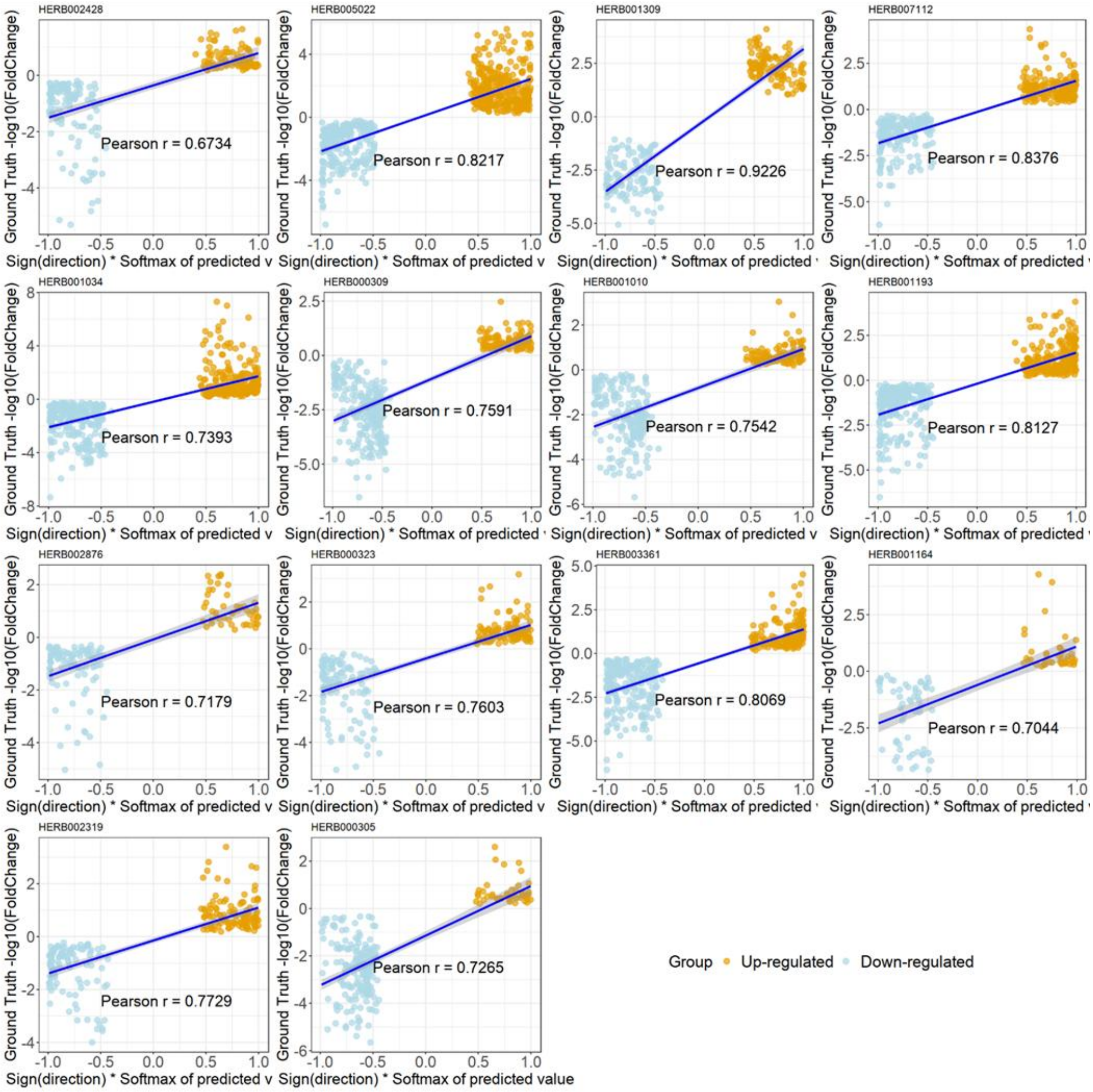

Supplementary Figure S15

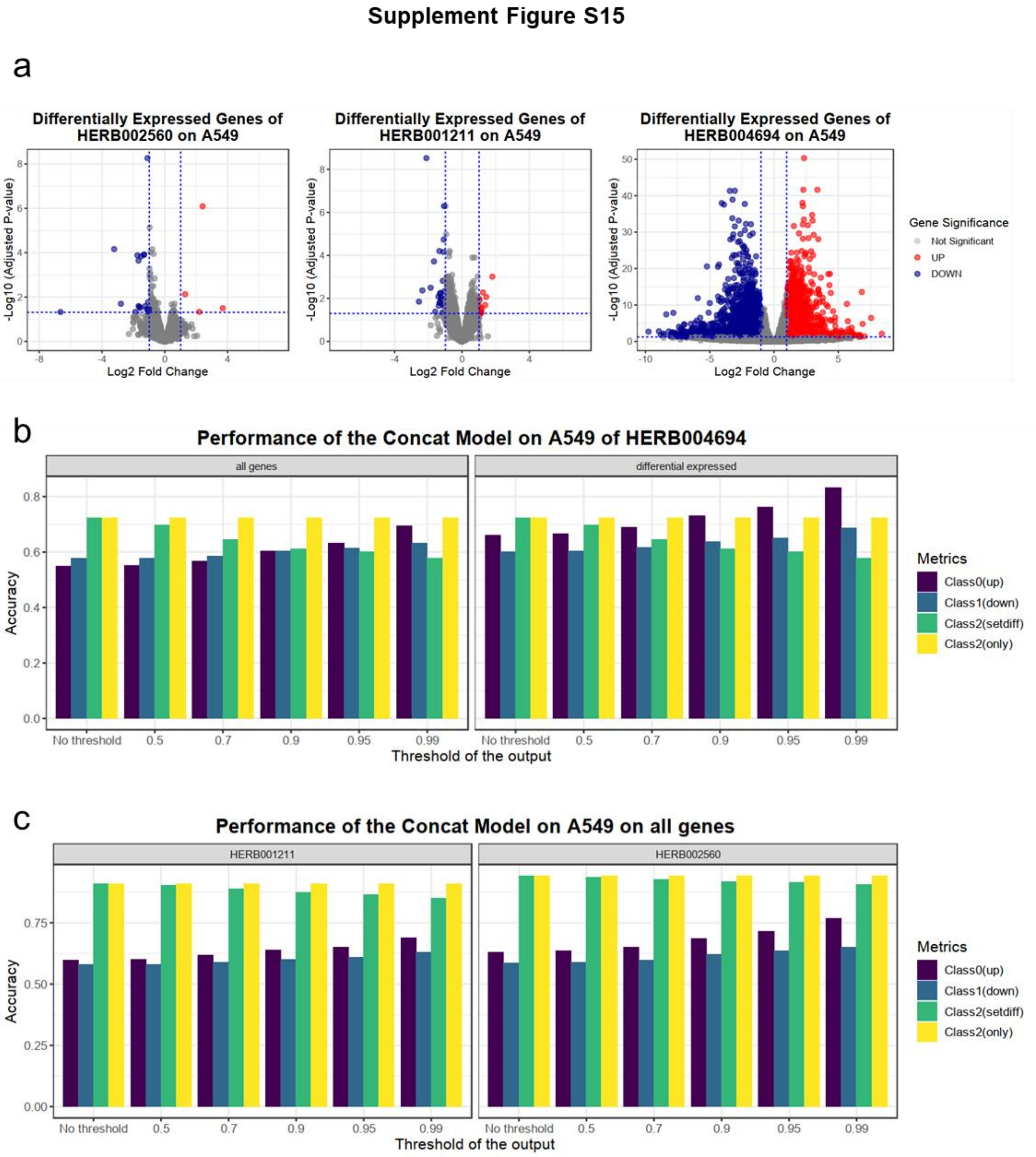

Supplementary Figure S16

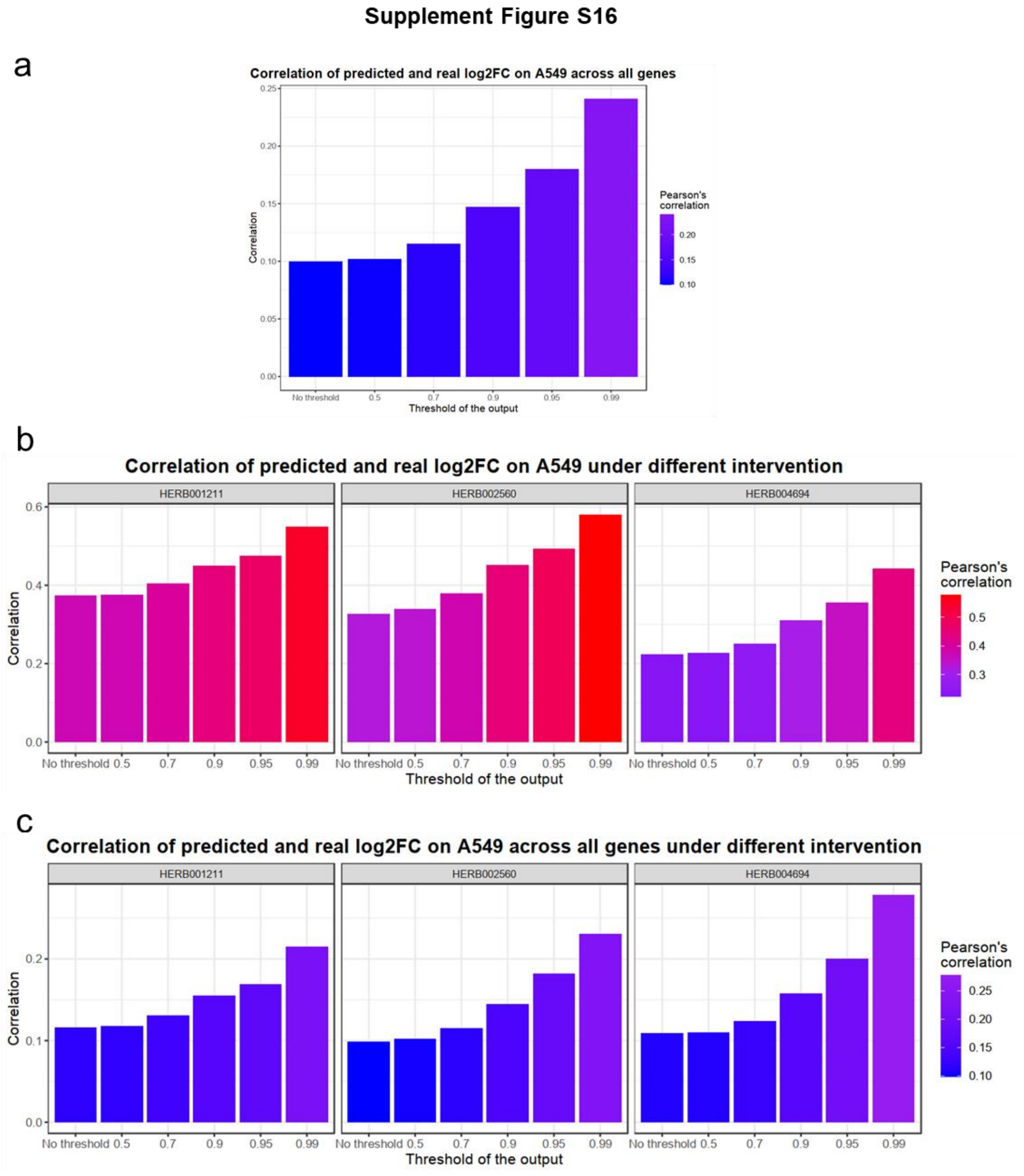

Supplementary Figure S17

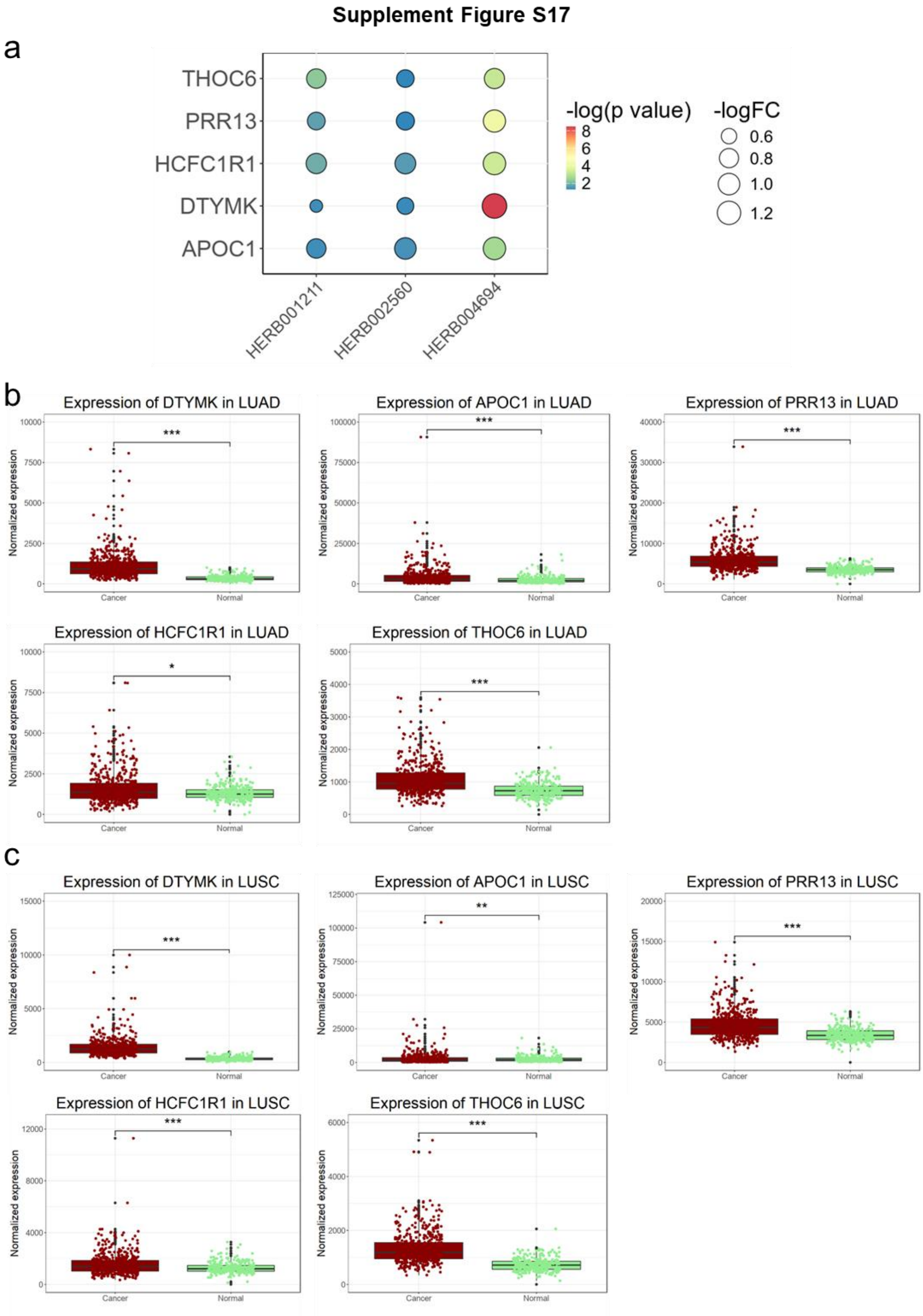

Supplementary Figure S18

**Supplement Figure S18**

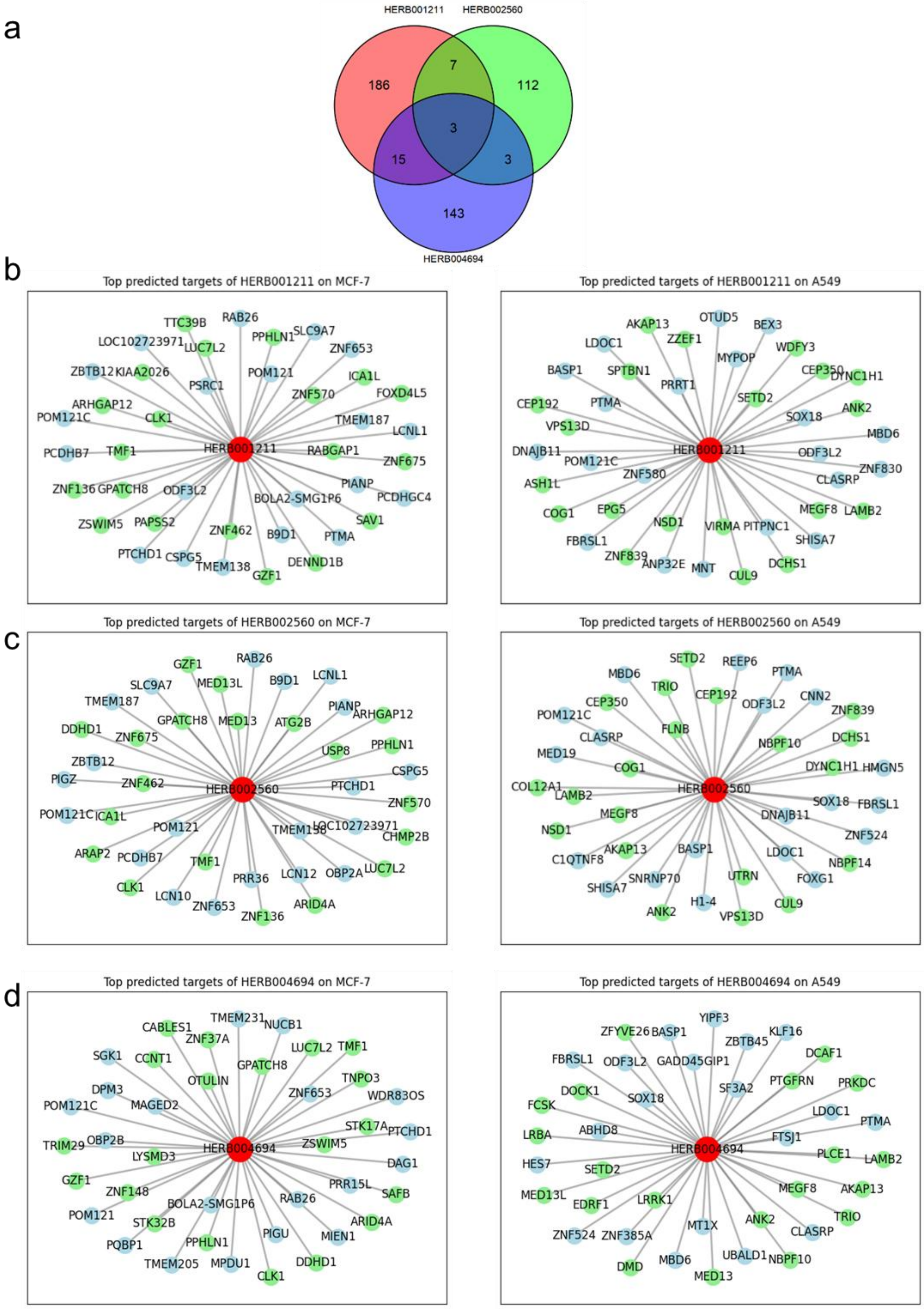

Supplementary Figure S19

**Supplement Figure S19**

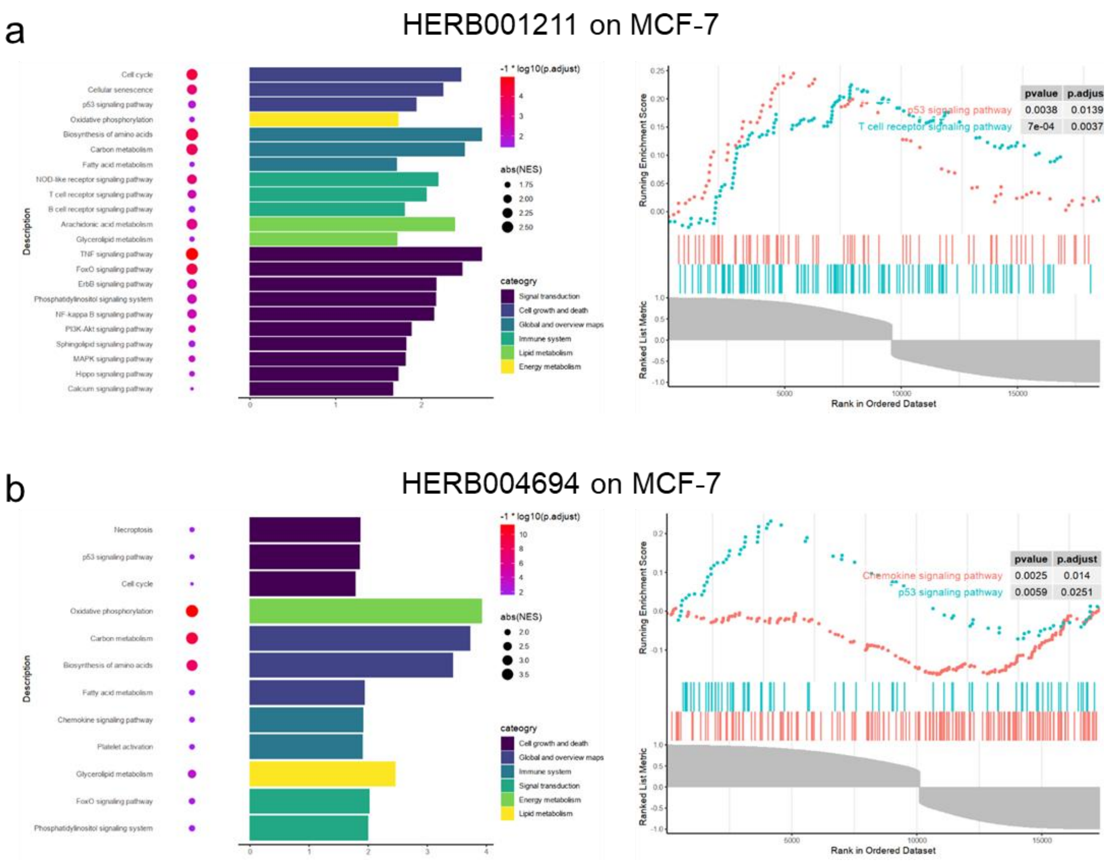

Supplementary Figure S20

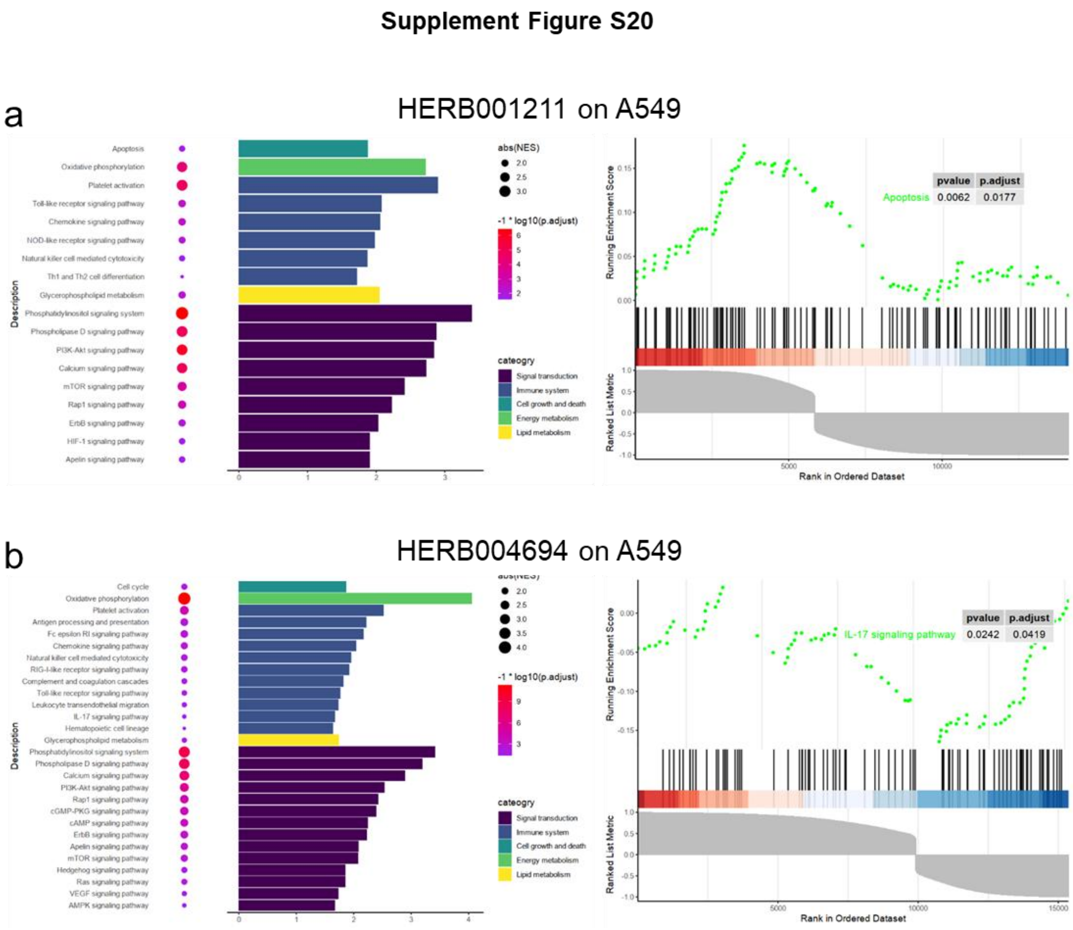

Supplementary Figure S21

**Supplement Figure S21**

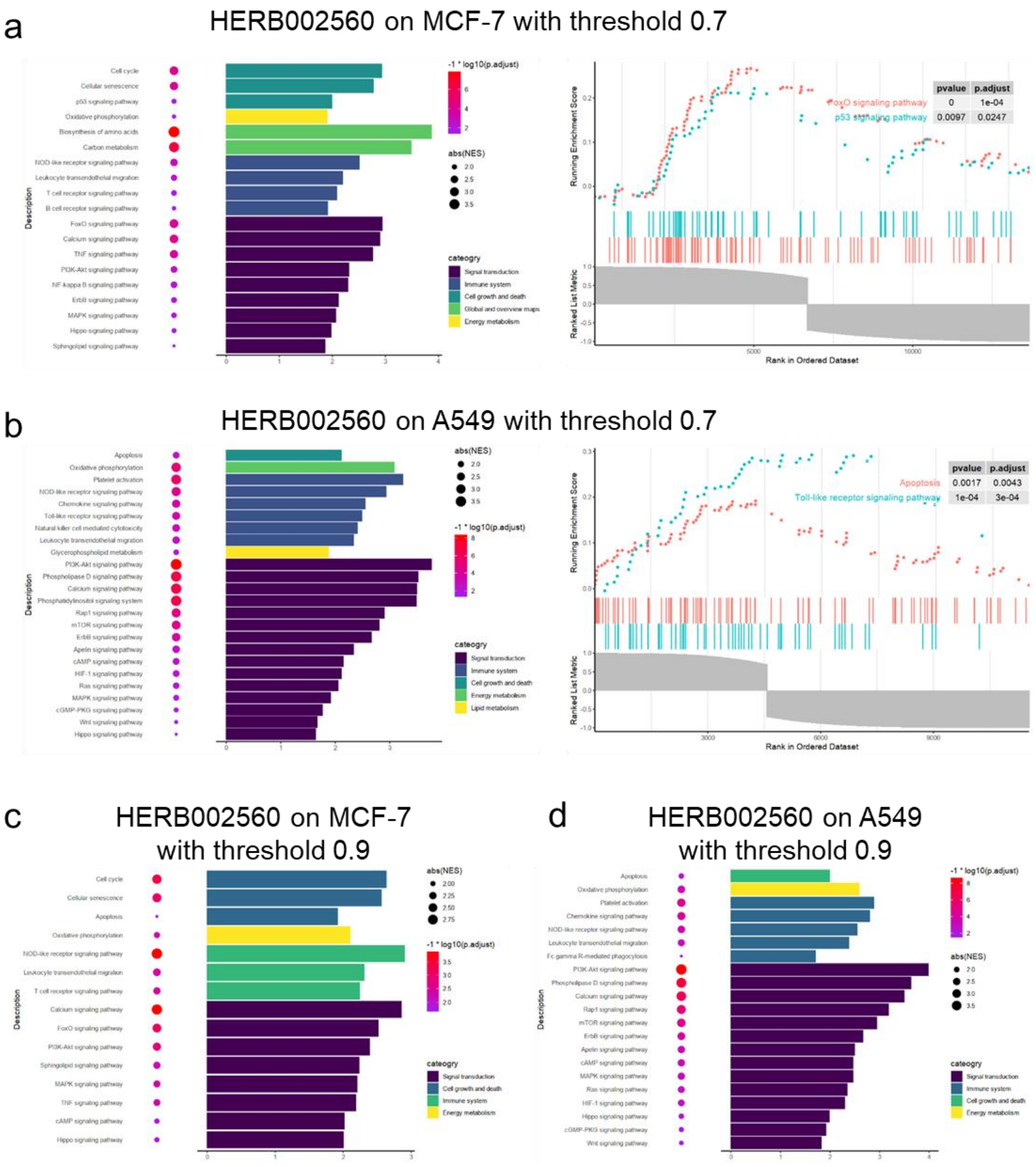

Supplementary Figure S22

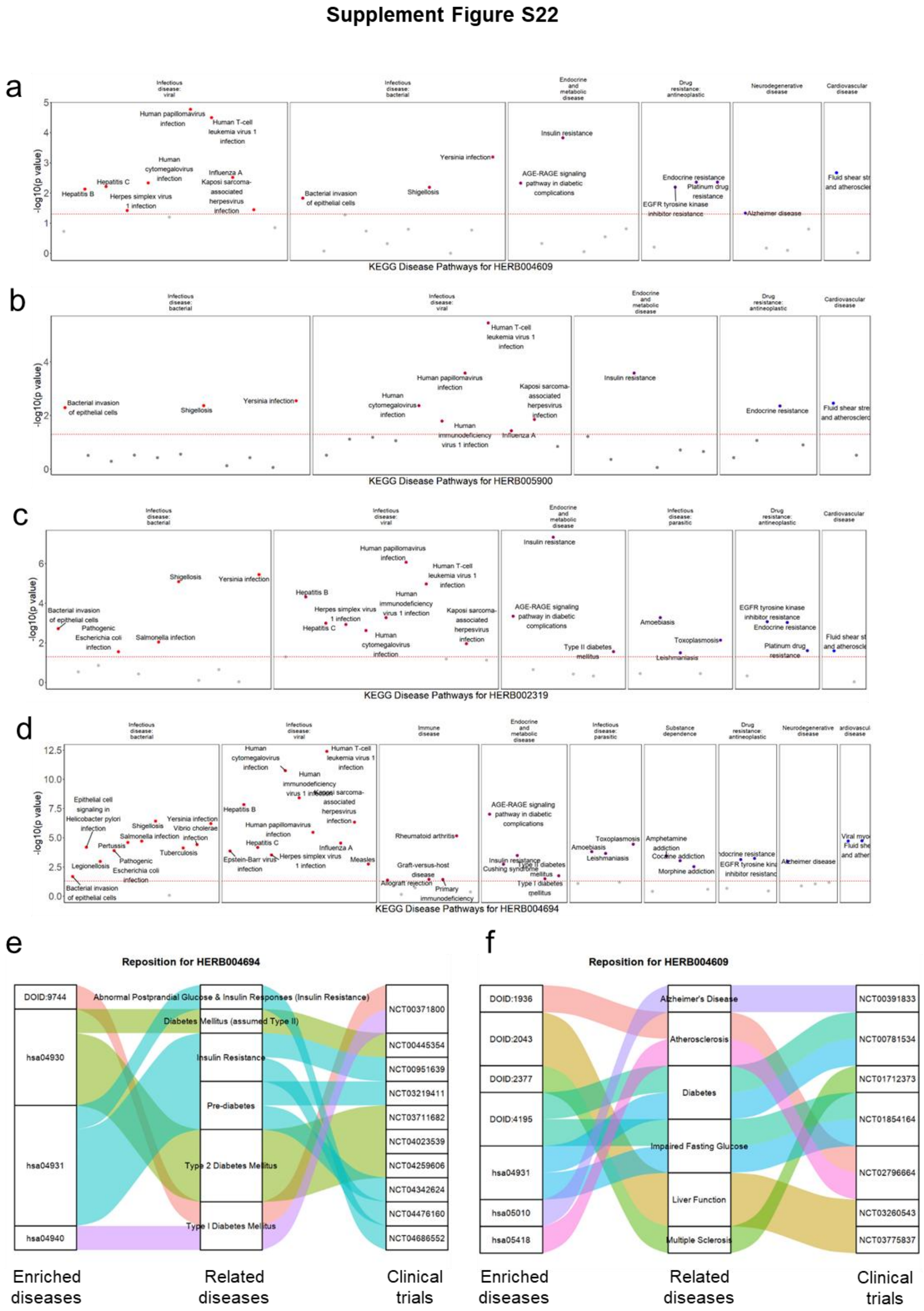

Supplementary Figure S23

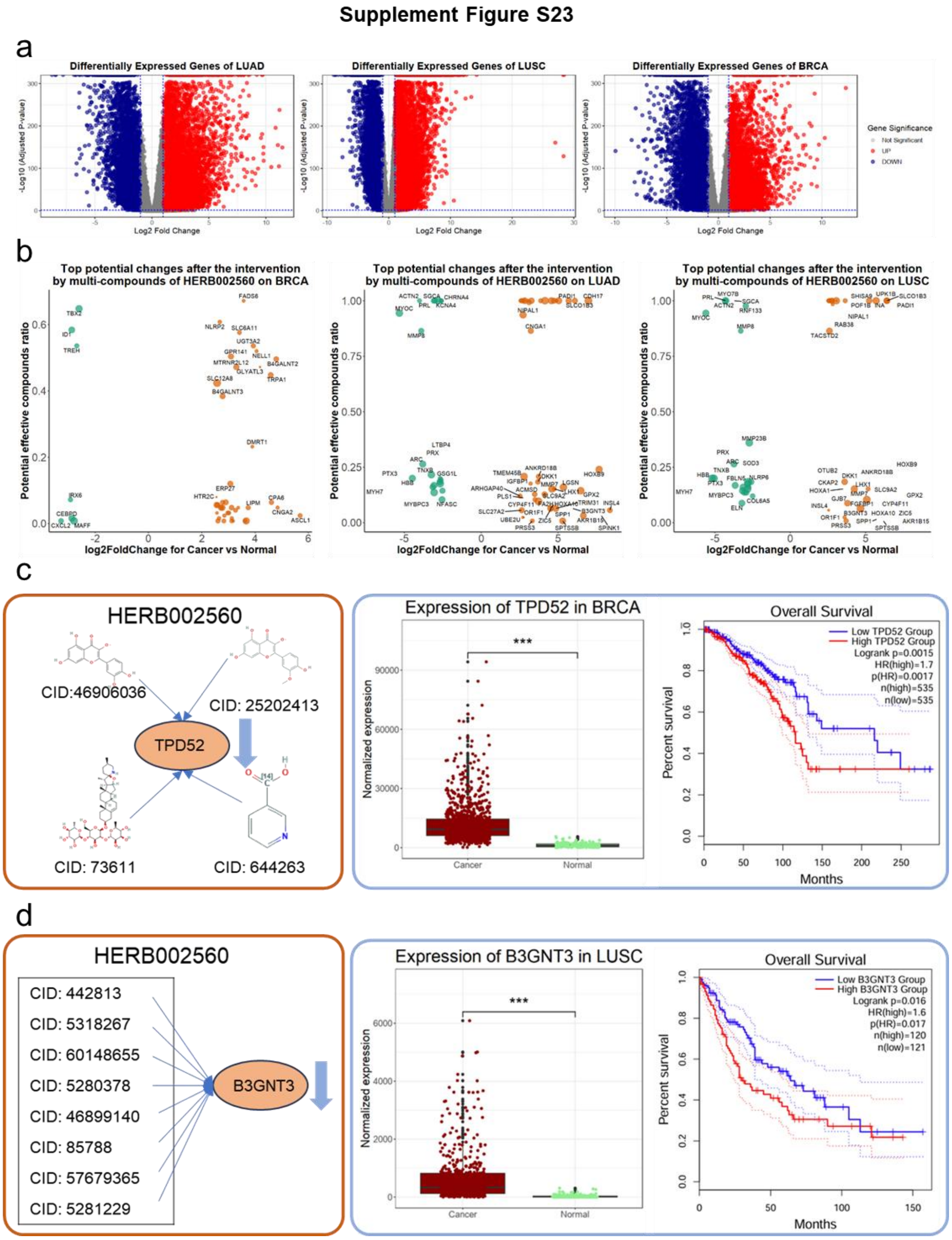

#### Supplementary Tables

Supplementary Table S1

| Parameters | Searching range |
|---|---|
| batch size | 1024, 512, 256, 128, 64 |
| learning rate | 1e-3, 1e-4, 1e-5, 1e-6 |
| L2 regularization | 1e-5, 1e-4, 1e-3, 1e-2 |
| dropout | 0.1, 0.2, 0.5, 0.8 |
| dimension in MLP layer | 512, 1024 |
| dimension in Set Transformer layer | 512, 1024, 2048 |

Supplementary Table S2

| Parameters | Searching range |
|---|---|
| batch size | 1024, 512, 256, 128, 64 |
| learning rate | 1e-3, 1e-4, 1e-5, 1e-6 |
| L2 regularization | 1e-5, 1e-4, 1e-3, 1e-2 |
| dropout | 0.1, 0.2, 0.5, 0.8 |

Supplementary Table S3

| epoch | train_loss | val_loss | val_acc | val_auc |
|---|---|---|---|---|
| 0 | 0.10441394 | 0.05324341 | 0.9812739 | 0.9987965 |
| 1 | 0.05155225 | 0.03958947 | 0.986271 | 0.9993037 |
| 2 | 0.04112046 | 0.03384728 | 0.9883974 | 0.9994766 |
| 3 | 0.03519382 | 0.02904077 | 0.9901183 | 0.9996113 |
| 4 | 0.03123626 | 0.02632332 | 0.9910326 | 0.999678 |
| 5 | 0.02828881 | 0.02437879 | 0.9917747 | 0.9997238 |
| 6 | 0.02604143 | 0.02251237 | 0.992409 | 0.9997612 |
| 7 | 0.02424367 | 0.02061381 | 0.9929859 | 0.9998 |
| 8 | 0.02271379 | 0.01918993 | 0.9934423 | 0.9998263 |
| 9 | 0.02131978 | 0.01939688 | 0.9935078 | 0.9998265 |

Supplementary Table S4

| epoch | train_loss | val_loss | val_acc | val_auc |
|---|---|---|---|---|
| 0 | 0.10701013 | 0.0540627 | 0.9811048 | 0.9987419 |
| 1 | 0.0520684 | 0.03944144 | 0.9862926 | 0.9993043 |
| 2 | 0.04086678 | 0.03285723 | 0.9886795 | 0.9995105 |
| 3 | 0.03480242 | 0.02901058 | 0.9901946 | 0.9996068 |
| 4 | 0.03088879 | 0.02569392 | 0.9912389 | 0.9996935 |
| 5 | 0.02794328 | 0.02301959 | 0.9921398 | 0.9997486 |
| 6 | 0.02564236 | 0.02184432 | 0.9925949 | 0.999774 |
| 7 | 0.02374276 | 0.02015439 | 0.9931734 | 0.9998074 |
| 8 | 0.02217382 | 0.01901249 | 0.9935783 | 0.999829 |
| 9 | 0.02084753 | 0.01827641 | 0.9938413 | 0.999837 |

Supplementary Table S5

| Methods | Parameters | Accuracy | F1 Score | AUC |
|---|---|---|---|---|
| NoSet | 96.11M | 0.8177 | 0.8228 | 0.9410 |
| MLP | 5.07M | **0.8581** | **0.8611** | **0.9661** |
| Concat | 199.75M | 0.8250 | 0.8299 | 0.9480 |
| Add | 173.01M | 0.8216 | 0.8266 | 0.9417 |

Supplementary Table S6

| Methods | Accuracy | F1 Score | AUC |
|---|---|---|---|
| KNN | 0.6401 | 0.6237 | 0.8067 |
| LDA | 0.5258 | 0.5297 | 0.7042 |
| DT | 0.7759 | 0.7763 | 0.8319 |
| NoSet(small) | **0.8517** | **0.8504** | **0.9581** |
| MLP(small) | 0.7769 | 0.7684 | 0.9203 |
| Concat(small) | 0.8495 | 0.8509 | 0.9564 |
| Add(small) | 0.8479 | 0.8495 | 0.9547 |

Supplementary Table S7

| Parameters | Searching range |
|---|---|
| batch size | 256, 128, 64 |
| learning rate | 1e-3, 1e-4, 1e-5, 1e-6 |
| L2 regularization | 1e-5, 1e-4, 1e-3, 1e-2 |
| dropout | 0.1, 0.2, 0.5, 0.8 |

Supplementary Table S8

| Methods | Accuracy | F1 Score | AUC |
|---|---|---|---|
| KNN(ft) | 0.7088 | 0.7097 | 0.8701 |
| LDA(ft) | 0.6627 | 0.6620 | 0.8325 |
| DT(ft) | 0.8804 | 0.8806 | 0.9105 |
| MLP(ft) | 0.8290 | 0.8316 | 0.9419 |
| MLP(pt+ft) | 0.8376 | 0.8399 | 0.9452 |
| NoSet(ft) | 0.8812 | 0.8821 | 0.9651 |
| NoSet(pt+ft) | 0.9065 | 0.9067 | 0.9711 |
| Add(ft) | 0.8737 | 0.8748 | 0.9615 |

| | | | |
|---|---|---|---|
| Add(pt+ft) | 0.9270 | 0.9272 | 0.9856 |
| Concat(ft) | 0.8758 | 0.8768 | 0.9630 |
| Concat(pt+ft) | **0.9386** | **0.9387** | **0.9888** |

Supplementary Table S9

| **Methods** | **Accuracy** | **F1 Score** | **AUC** |
|---|---|---|---|
| Add(ft) | 0.8737 | 0.8748 | 0.9615 |
| Add(pt) | 0.3386 | 0.2323 | 0.5063 |
| Add(pt+ft) | **0.9270** | **0.9272** | **0.9856** |
| Concat(ft) | 0.8758 | 0.8768 | 0.9630 |
| Concat(pt) | 0.3379 | 0.2308 | 0.5162 |
| Concat(pt+ft) | **0.9386** | **0.9387** | **0.9888** |

Supplementary Table S10

| **Methods** | **Accuracy** | **F1 Score** | **AUC** |
|---|---|---|---|
| KNN | 0.6945 | 0.6948 | 0.8623 |
| DT | 0.8633 | 0.8633 | 0.8976 |
| LDA | 0.4393 | 0.4412 | 0.6174 |
| MLP(small pt+ft) | 0.8766 | 0.8778 | 0.9623 |
| NoSet(small pt+ft) | 0.8818 | 0.8827 | 0.9649 |
| Add(small pt+ft) | 0.8781 | 0.8791 | 0.9621 |
| Add(pt+ft) | 0.9270 | 0.9272 | 0.9856 |
| Concat(small pt+ft) | 0.8737 | 0.8748 | 0.9612 |
| Concat(pt+ft) | **0.9386** | **0.9387** | **0.9888** |

Supplementary Table S11

| **Methods** | **Accuracy** | **F1 Score** | **AUC** |
|---|---|---|---|
| KNN | 0.7062 | 0.7108 | 0.8628 |
| DT | 0.6966 | 0.6919 | 0.7725 |
| LDA | 0.5792 | 0.5776 | 0.6945 |
| Vanilla NN | 0.7816 | 0.7847 | 0.9161 |

| | | | |
|---|---|---|---|
| Add | 0.8262 | 0.8288 | **0.9373** |
| Concat | **0.8275** | **0.8299** | 0.9365 |
